# Evaluation of an Instrumented Helmet for Measurement of Orthosis–Head Interface Forces in Pediatric Craniosynostosis

**DOI:** 10.64898/2026.11.20.767636

**Authors:** Ahmet Arif Colakoglu, Yunus Samiepour, Ce Liang, Anastasiia Maliuk, Damith Chathuranga, Hans Delye, Mehran Moazen, Peter Culmer, Ali Alazmani

## Abstract

Postoperative cranial orthoses guide skull growth after endoscopy-assisted craniosynostosis surgery in infants, yet the forces they apply at the orthosis–head interface remain poorly characterized. Here, a cranial helmet with 12 soft magnetic tactile sensors is developed to measure these forces. The helmet, a computed tomography (CT)-derived replica of a patient’s first clinical helmet, is evaluated on patient-derived expandable head phantoms with and without a modeled craniotomy. The phantoms follow the first treatment phase of a female infant with unicoronal craniosynostosis, from surgery at 77 days to the end of the first helmet period at 160 days (843 mL), extended to 175 days (860 mL). As the phantom expands, contact develops progressively and remains non-uniform, with occipital and spheno-temporal regions as the dominant load-bearing sites. At 175 days, mean local forces reach 258.3 ± 8.7 gf occipitally, 168.5 ± 5.9 gf and 104.5 ± 39.9 gf at two spheno-temporal sites, and 105.7 ± 13.6 gf frontally. Repeated helmet fitting reproduces the sensor-level loading pattern. Compared with the phantom without the helmet, CT shows reduced peripheral displacement and redirection of expansion toward superior relief regions. The instrumented helmet described here provides a quantitative testbed for orthosis design and computational-model validation.

## 1 Introduction

Craniosynostosis is the premature fusion of one or more cranial sutures and affects approximately 1 in 2,000– 2,500 live births [1–3]. Unicoronal craniosynostosis occurs in about 1 in 10,000 live births and is more common in females [4]. In this form, premature fusion of one coronal suture alters skull growth, producing an asymmetric forehead, flattened on the fused side and bulging on the opposite side (anterior plagiocephaly) [5]. This form is also associated with ophthalmic complications, most commonly strabismus and astigmatism [6]. For infants treated early, minimally invasive endoscopic surgery to release the fused suture may be followed by several months of cranial orthosis therapy to guide skull growth toward a more typical morphology [7,8].

Cranial orthoses act by constraining growth in selected regions while allowing expansion into designed relief spaces [7–9]. However, the forces generated at the orthosis–head interface remain poorly characterized. Computational models have simulated helmet therapy and predicted the resulting intracranial contact pressures and sutural stresses [10,11], and related finite-element frameworks predict skull growth after different surgical reconstructions [12,13]. These predictions have not yet been validated against direct physical measurements, so their absolute values should be interpreted with caution [10]. Force sensing has been applied at related interfaces: sensor arrays embedded in protective helmets showed highly uneven head– helmet contact forces associated with head shape [14]. These helmet measurements, however, were obtained in adult volunteers under static fit rather than during prolonged orthotic loading of a growing infant head. Direct evidence for cranial orthoses is very limited: head–orthosis pressures have so far been measured *in vitro* only on static, rigid head molds of infants with positional plagiocephaly (head flattening caused by the infant’s lying position, without suture fusion) [15], and a handheld instrumented ruler for checking helmet fit has been tested only in bench compression tests [16]. Beyond the head, the soft magnetic tactile sensors on which the present sensor design is based have measured the loads transmitted across a hand splint–skin interface during activities of daily living [17]. Animal models differ from human infant cranial anatomy in geometry and suture architecture [18,19], and static skull phantoms reproduce anatomy but not progressive cranial expansion [20,21].

There is therefore a need for an instrumented cranial orthosis capable of directly measuring local interface forces during controlled head expansion. Such measurements could establish the magnitude and spatial distribution of orthotic loading, provide experimental data for computational-model validation, and support more quantitative assessment of different orthosis designs.

The present work builds on a previously developed *in vitro* platform in which CT-derived infant head phantoms, consisting of printed cranial bone plates joined by latex sutures and covered by latex layers around a fluid-filled intracranial cavity, are expanded in controlled volume steps by a pneumatic–hydraulic system to follow age-specific intracranial volume targets [22]. That platform tracked head shape as the phantom expanded but provided no information on the loads applied by an orthosis. The focus here is the second element: a replica of the patient’s clinical helmet with soft magnetic tactile force sensors integrated in its shell, which measures local contact forces at the orthosis–head interface. Combining the two (Figure 1) allows a clinical helmet to be loaded by a head that expands in a known and repeatable way, so that the onset, magnitude and distribution of contact can be measured under conditions that cannot be controlled or instrumented in an infant.

**Figure 1.**
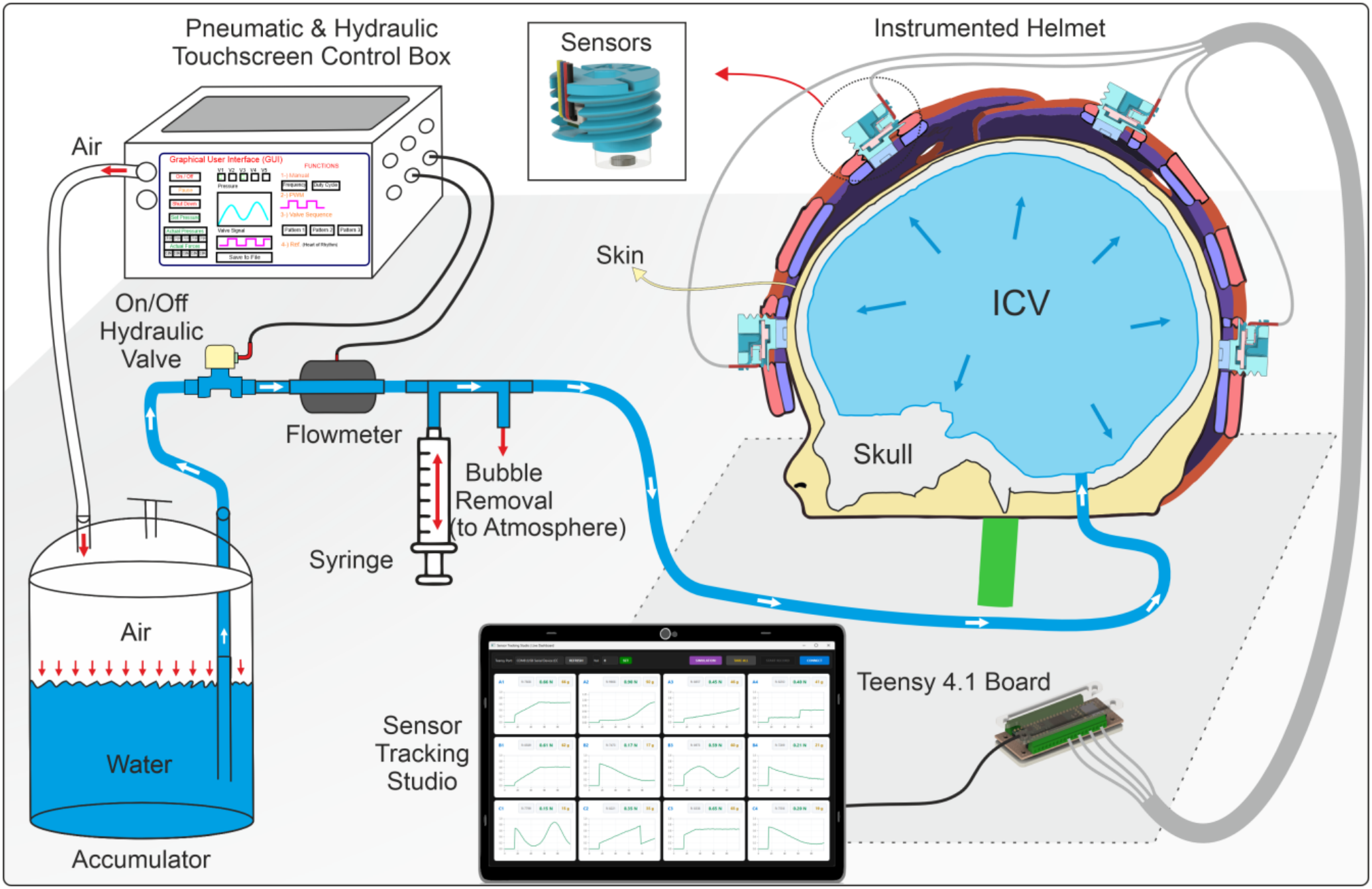
Concept of the *in vitro* platform. Left: pneumatic–hydraulic expansion system; a touchscreen control box pressurizes the water accumulator, and water passes through an on/off hydraulic valve and a flowmeter to the phantom, with a syringe branch for bubble removal. Right: CT-derived head phantom (skull, intracranial volume (ICV) and skin), which expands outward as fluid is delivered (blue arrows), fitted with the instrumented helmet, a full-scale replica of the patient’s first clinical helmet carrying 12 soft magnetic tactile force sensors (inset: one sensor). Bottom: the sensors are read by a Teensy 4.1 microcontroller, and the 12 channels are displayed and recorded in real time in Sensor Tracking Studio.

Here, a computed tomography (CT)-derived replica of a patient’s first clinical helmet was developed and instrumented with 12 soft magnetic tactile force sensors (Figure 1). The helmet was evaluated on patient-derived expandable head phantoms following the first of three helmet-treatment phases of a female infant with unicoronal craniosynostosis (Figure 2), from surgery at 77 days to the end of Helmet 1 at 160 days (843 mL), extended to 175 days (860 mL). This setup was used to (i) quantify the magnitude and spatial distribution of local orthosis–head interface forces during cranial expansion, (ii) assess the repeatability of the measured force pattern following helmet removal and refitting, and (iii) determine how the helmet alters cranial surface displacement relative to the same phantom without the helmet.

**Figure 2.**
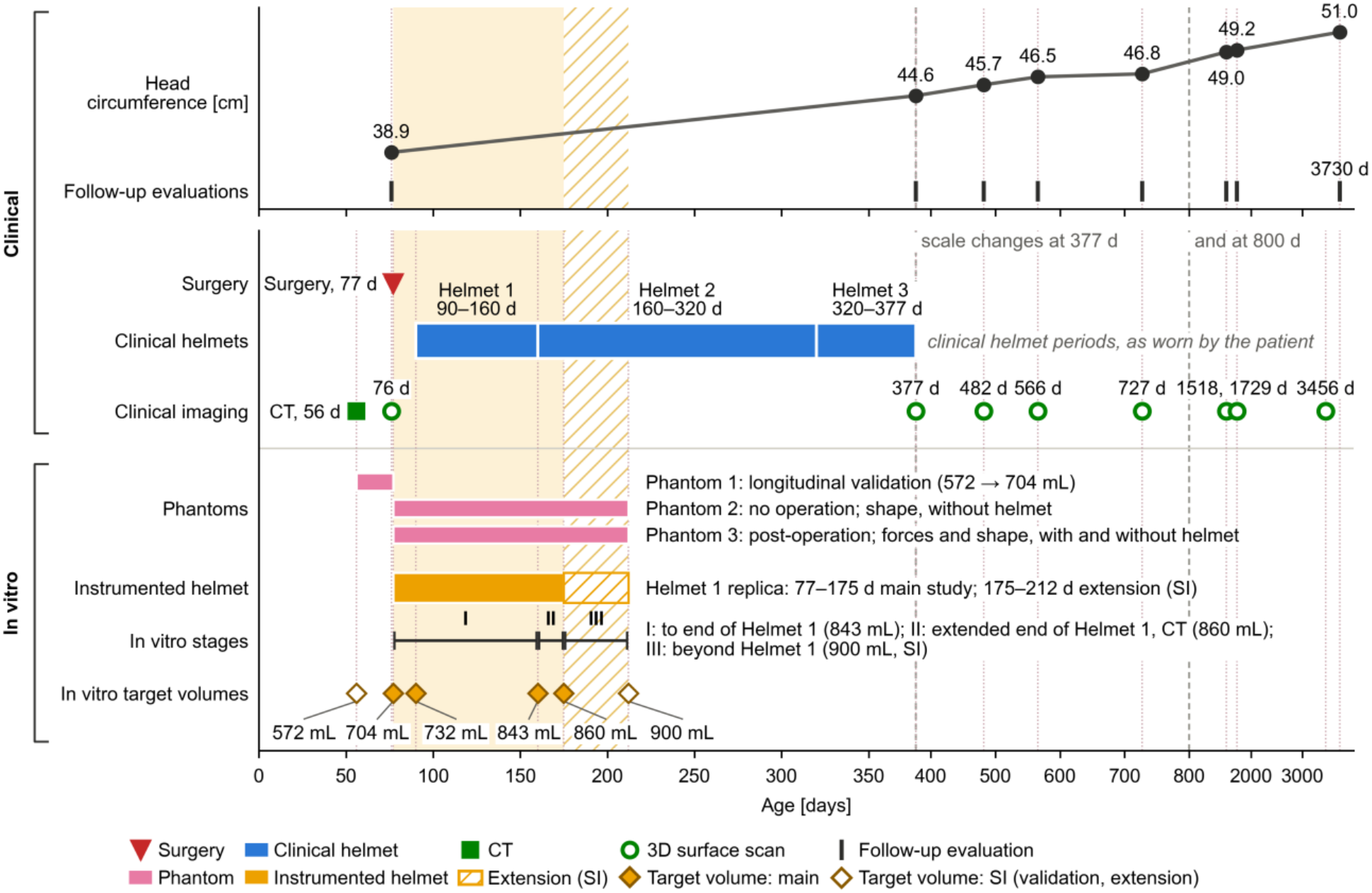
Clinical treatment period (56–3730 days) and *in vitro* study design. Clinical: head circumference measured at the follow-up evaluations (76–3730 days); surgery at 77 days; the three clinical helmet periods as worn by the patient (Helmet 1, 90–160 days; Helmet 2, 160–320 days; Helmet 3, 320–377 days); and clinical imaging (CT of the skull and skin at 56 days; 3D surface scans of the skin from 76 to 3456 days). *In vitro*: the three phantoms and their roles; the instrumented helmet (a replica of Helmet 1), fitted from 77 to 175 days in the main study and to 212 days in the extension (Sections S7 and S8, Supporting Information); *in vitro* stages I–III; and the target volumes from the normative ICV curve (filled diamonds, main study; open diamonds, Supporting Information). The shaded band marks the main *in vitro* study period (77–175 days) and the hatched band the extension (175–212 days). The time axis is piecewise linear, with scale changes at 377 and 800 days.

## 2 Results

### 2.1 Instrumented Helmet and Head Phantoms

The instrumented helmet is a full-scale (1:1) copy of the patient’s first clinical helmet (Helmet 1), reproduced from CT and fitted with 12 soft magnetic tactile force sensors in three regions: frontal (A1–A4), occipital (B1– B4) and spheno-temporal (C1–C2 left, C3–C4 right) (Figures 1 and 3). For CT imaging, a metal-free dummy helmet with the same geometry was used.

**Figure 3.**
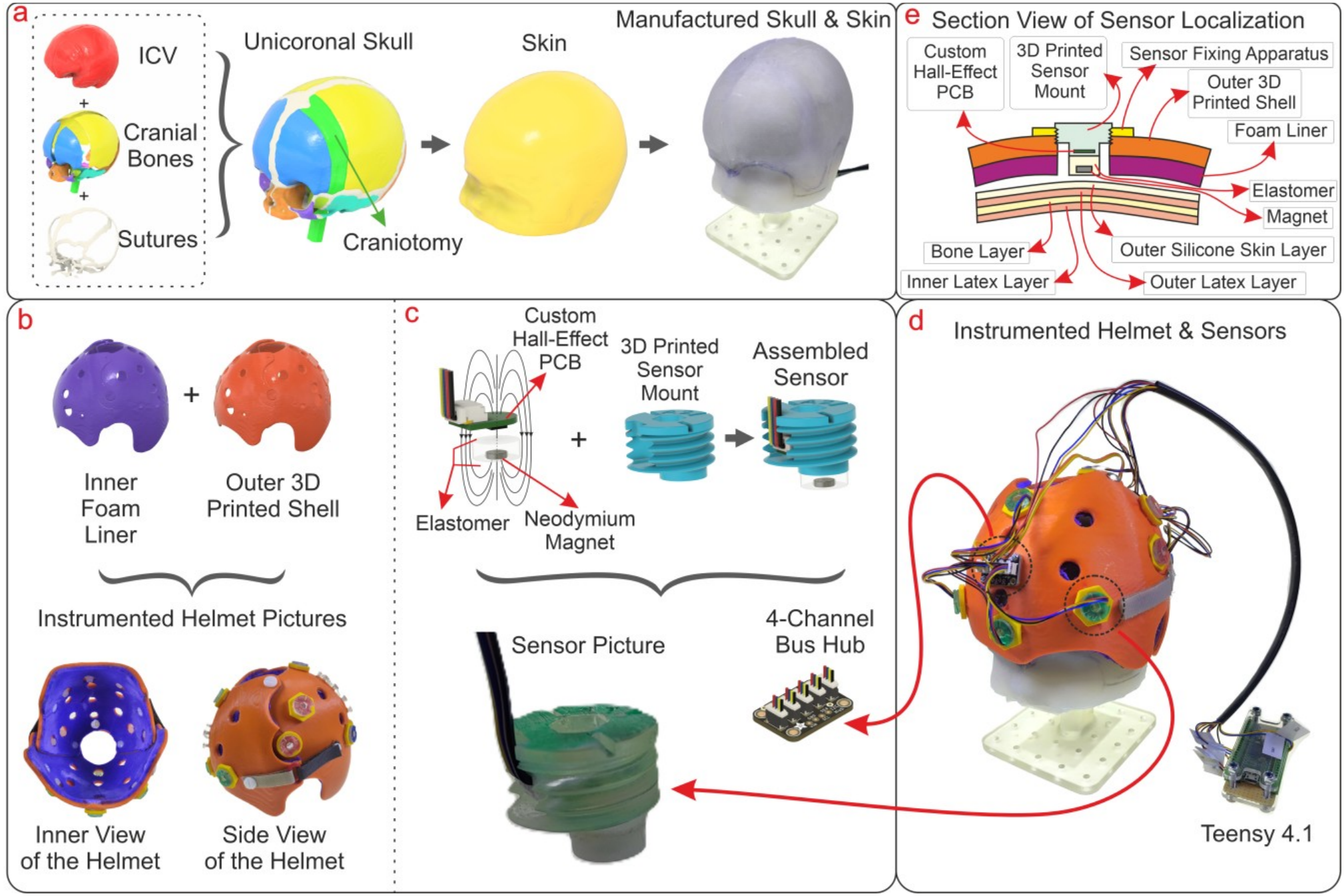
Phantom and instrumented helmet. (a) Phantom assembly: cranial bone plates and latex sutures around a sacrificial ICV core, covered by a silicone skin. (b) ABS helmet shell and foam liner. (c) Assembled views of one force sensor, with a bus hub. (d) Integrated helmeted phantom with wiring and microcontroller. (e) Section through a mounted sensor at the interface.

Three patient-derived head phantoms were built and are numbered in the order of the modeling sequence (Section S5, Table S3, Supporting Information). Phantom 1 was made directly from the patient’s 56-day CT (572 mL) and was expanded to 77 days (704 mL) only to validate the expansion against the patient’s surface scan (longitudinal validation; Section S8, Figure S14, Supporting Information). Phantoms 2 and 3 were made from the 77-day surgery-day geometry predicted by the companion finite-element growth model: in Phantom 2 (no operation), the fused coronal suture was kept as one rigid part; in Phantom 3 (post-operation), a surgeon-defined craniotomy was modeled across it. Phantoms 2 and 3 were expanded from 77 days (704 mL) in controlled volume steps that translate clinical time points into intracranial volume (ICV): 90 days (732 mL, clinical fitting of Helmet 1), 160 days (843 mL, end of Helmet 1) and 175 days (860 mL, an extended end-of-Helmet-1 step at which *in vitro* CT was acquired; Section 5.2). The instrumented helmet was fitted from the 77-day baseline (Section 5.2), and all force measurements were made on Phantom 3 over seven independent helmet fittings (n = 7).

### 2.2 Phantom Validation

In a baseline validation, the external surfaces of Phantoms 2 and 3 at 77 days (704 mL) were compared with the patient’s pre-operative surface scan at 76 days (Figure 4). The 1-day difference corresponds to about 2 mL of intracranial volume on the target-volume curve (701.8 vs 704.0 mL, 0.3%) and was considered negligible; 77 days (704 mL) was used because it is the surgery-day baseline of the finite-element geometry. On the common color scale, most of the cranial vault fell within 0–2.5 mm, with limited local regions approaching 5 mm. The differences along the lower edge coincided with missing coverage in the clinical scan and were treated as acquisition gaps. A longitudinal validation, in which Phantom 1 was expanded from 56 to 77 days (572 to 704 mL) and compared with the same scan, is reported in Section S8, Figure S14 (Supporting Information).

**Figure 4.**
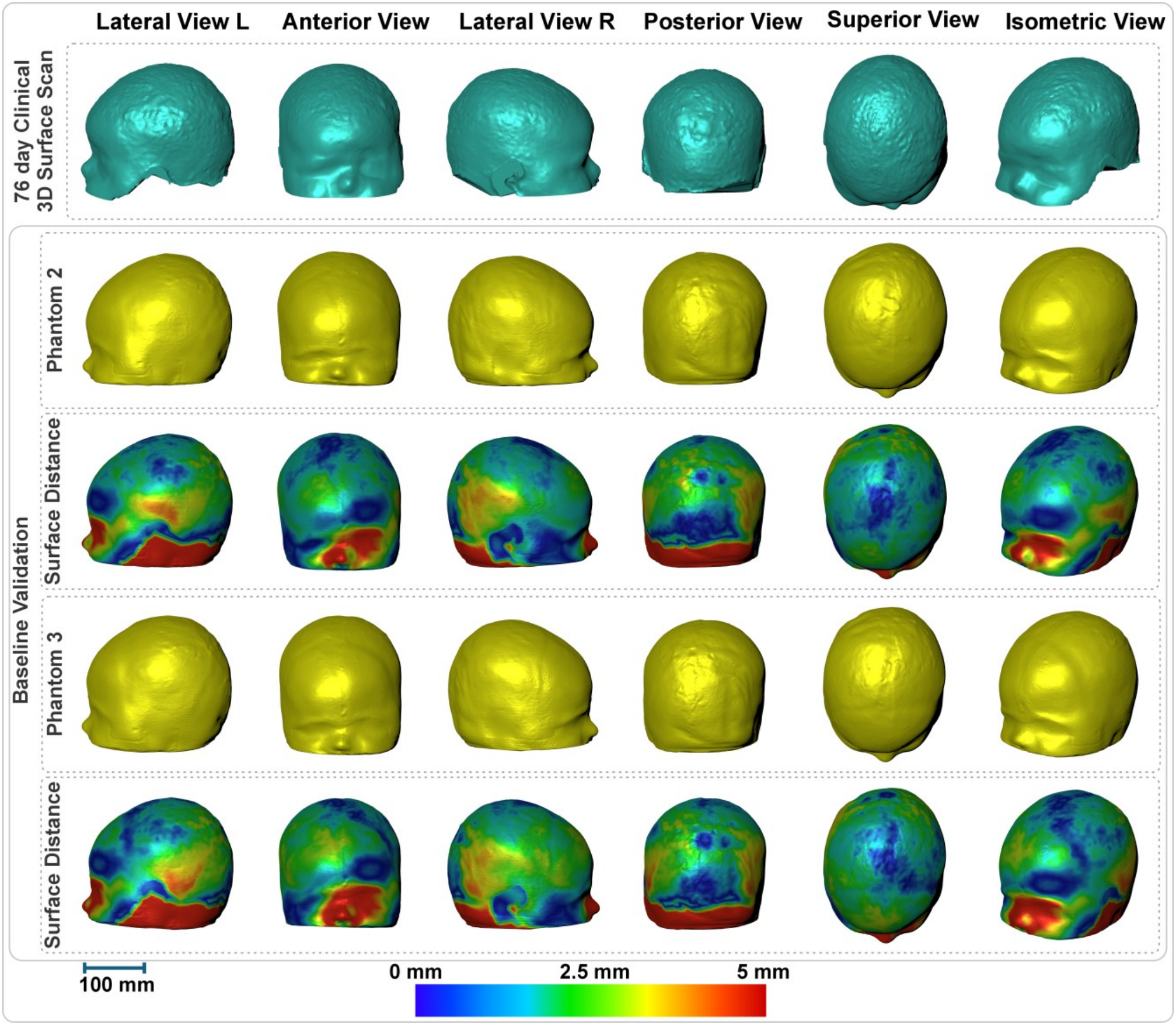
Baseline validation: comparison of Phantoms 2 and 3 at 77 days (704 mL) with the patient’s 76-day pre-operative surface scan. All surface-distance maps share the same 0–5-mm color scale.

### 2.3 Local Contact Forces and Fitting Repeatability

The representative continuous recording showed that contact developed sequentially rather than uniformly across the helmet–head interface (Figure 5a). Sensors already engaged at 77 days (704 mL) generally remained load bearing and increased as intracranial volume rose. Sensors located within initially relieved regions remained at or near zero until later in the expansion, after which positive force developed as the external surface approached and engaged the liner.

**Figure 5.**
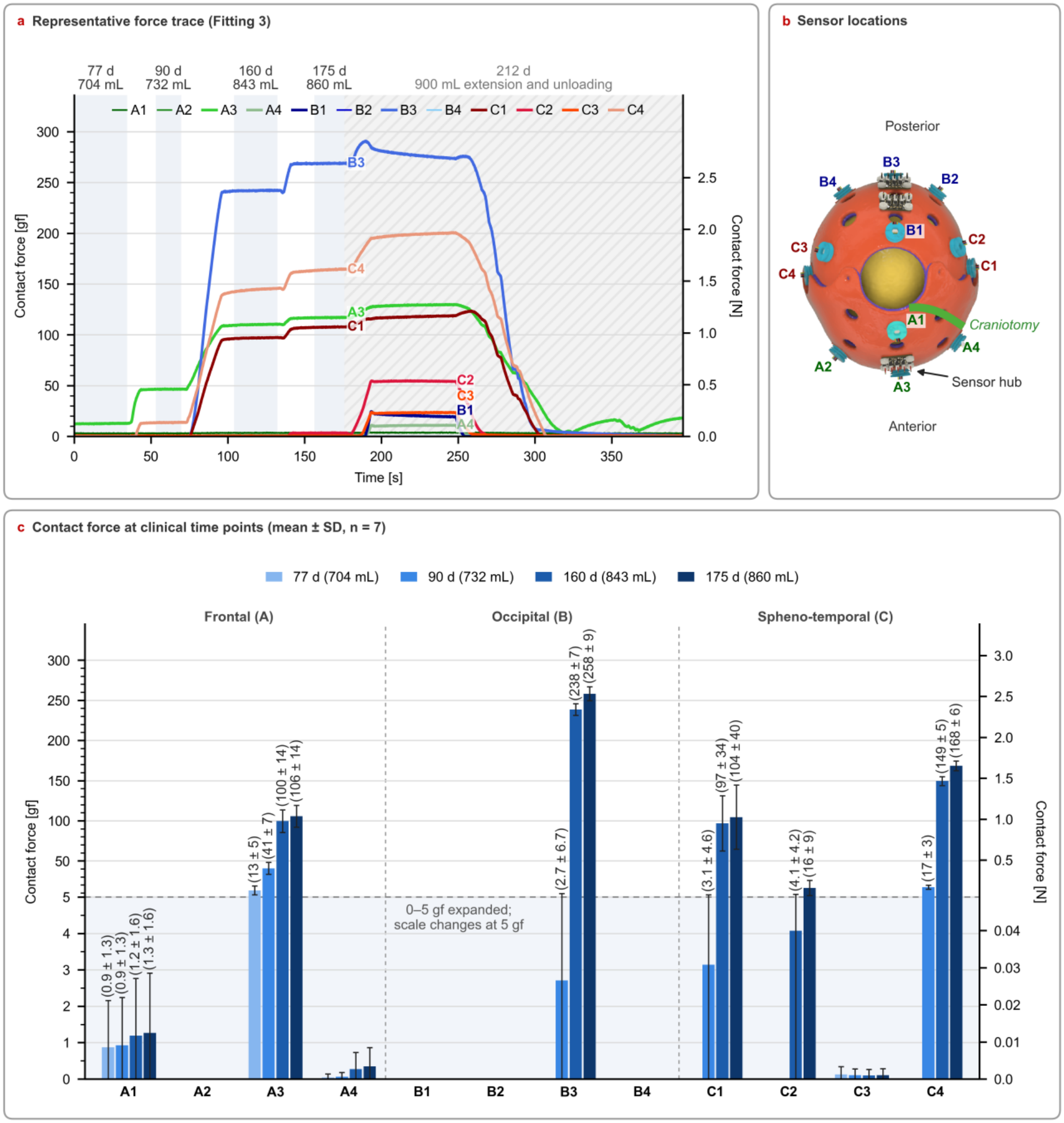
Interface forces on Phantom 3 (modeled post-craniotomy). (a) Force trace of a representative fitting (Fitting 3, closest to the series mean); shaded bands mark stable holds, and the hatched 212-day (900-mL) extension is reported in Section S7, Figure S12 (Supporting Information). (b) Sensor sites: A frontal, B occipital, C spheno-temporal (C1–C2 left, C3–C4 right). (c) Mean ± SD force at each volume step over seven fittings (n = 7); the force axis is expanded between 0 and 5 gf, with a change of scale at 5 gf; sensors without bars recorded no measurable contact (< 0.5 gf).

At 77 days (704 mL), only the frontal site A3 carried measurable load (13.3 ± 5.4 gf); all other sensors recorded less than 1 gf (Figure 5c). At the expanded intracranial volume corresponding to the clinical fitting of Helmet 1 at 90 days (732 mL), A3 had increased to 40.8 ± 7.3 gf and the right spheno-temporal site C4 had engaged (17.3 ± 2.6 gf), with initial contact at C1 and B3 in some fittings. By the end of Helmet 1 at 160 days (843 mL), load had spread to the occipital and spheno-temporal regions: B3 reached 238.5 ± 7.2 gf, C4 149.4 ± 5.4 gf, C1 96.7 ± 34.4 gf and A3 99.5 ± 14.1 gf.

At 175 days (860 mL), force remained strongly non-uniform. B3 carried the highest mean force, 258.3 ± 8.7 gf, followed by C4 at 168.5 ± 5.9 gf, A3 at 105.7 ± 13.6 gf and C1 at 104.5 ± 39.9 gf, with C2 at 16.3 ± 9.4 gf. The highest single-site force was 269 gf, at B3, corresponding to a nominal sensor-face pressure of approximately 52 kPa.

Regional load shares changed as the head came into contact with the helmet. At 77 days (704 mL), the frontal group accounted for 99% of the summed sensor load; at 160 and 175 days (843 and 860 mL), the occipital group (41% and 40%) and the spheno-temporal group (42% and 44%) together carried 83–84%, with the frontal group at 16–17%. Sites A2, B1, B2, B4 and C3 recorded no measurable force up to 175 days (860 mL), although their raw signals remained active throughout; B1 and C3 engaged only in the 212-day (900-mL) extension (Section S7, Table S5, Supporting Information).

Removing and refitting the helmet reproduced the sensor-level force pattern: the same dominant and unloaded sites were retained in all seven fittings, and at 175 days (860 mL) the standard deviation was 3–4% of the mean at B3 and C4 and 13% at A3. The largest variation occurred at C1 (coefficient of variation 38%), and the force at this site decreased progressively over the series, from 168.5 gf in the first fitting to 55–57 gf in the last two; A3 was also lower in the last two fittings (Section S7, Figure S12 and Table S6, Supporting Information). The maximum between-fitting range in regional load share at 175 days was 9 percentage points (occipital group, 35–44%).

Complete sensor-level values at every volume step, including the 212-day (900-mL) extension, and the continuous traces of all seven fittings are provided in Section S7, Figure S12 and Tables S5–S7 (Supporting Information).

### 2.4 Helmet-Associated Changes in Cranial Morphology

Comparison of Phantoms 2 and 3 showed the effect of releasing the fused left coronal suture (Figure 6). Phantom 2 showed a localized region of high displacement on the right side of the skull, opposite the fused left coronal suture. In Phantom 3, this right-sided region was smaller and expansion increased around the modeled release, as seen in the lateral and isometric views.

**Figure 6.**
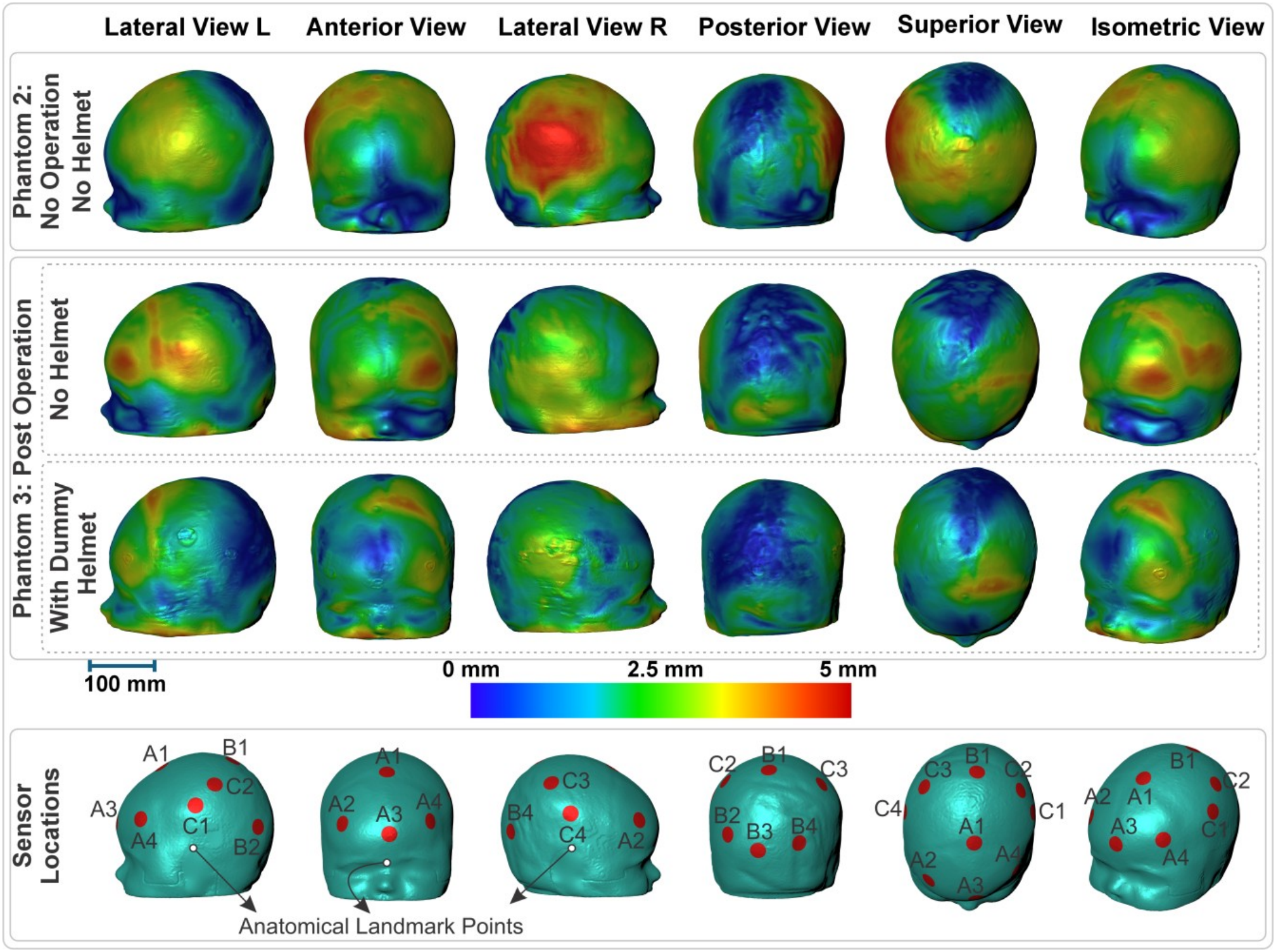
Cranial surface displacement from 77 days (704 mL) to 175 days (860 mL) for three configurations: Phantom 2 without a helmet, Phantom 3 without a helmet and Phantom 3 with the dummy helmet. Bottom row: approximate projected sensor positions, shown for anatomical reference only (not measured coordinates).

The most direct assessment of the helmet effect was the within-build comparison of Phantom 3 with and without the helmet (Figure 6). Without the helmet, the modeled post-craniotomy phantom showed pronounced outward displacement around the released region and adjacent lateral vault. With the helmet fitted, the regions showing the greatest lateral, frontal and occipital displacement in the no-helmet condition were visually reduced to intermediate levels on the common color scale, while expansion became more prominent toward the superior vault. Maps of the 212-day (900-mL) extension for the same three configurations are provided in Section S8, Figure S13 (Supporting Information).

The section overlays provided the geometric basis for this shift toward the superior vault (Figure 7). At 77 days (704 mL), the sagittal and coronal planes showed the largest clearance between the cranial surface and helmet over the superior vault. By 175 days (860 mL), the expanded surface had entered and largely filled these spaces, approaching the liner. In the horizontal plane, the frontal, occipital and lateral surfaces were already close to or in contact with the helmet at baseline and moved outward comparatively less. The fact that this clearance was largely filled at 175 days (860 mL), shortly after the clinical end of Helmet 1 at 160 days (843 mL), is consistent with the timing of the clinical transition to Helmet 2 in this experimental model.

**Figure 7.**
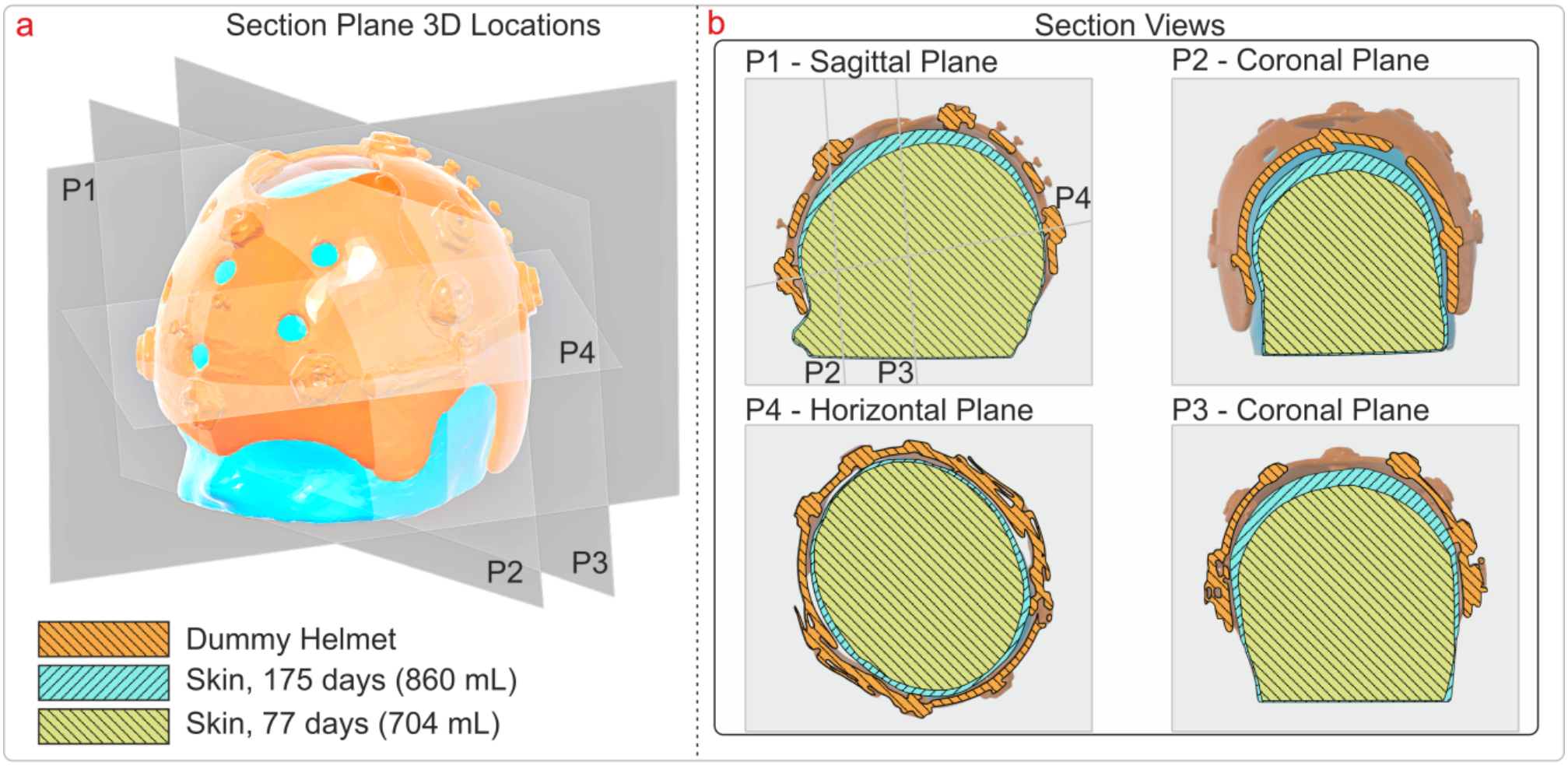
Helmet clearance during expansion of Phantom 3. (a) Locations of section planes P1–P4. (b) Overlaid sections of the dummy helmet and of the skin at 77 and 175 days (704 and 860 mL).

## 3 Discussion

To the best of our knowledge, this is the first instrumented replica of a clinical helmet used to measure orthosis– head interface forces during controlled expansion of a patient-derived craniosynostosis head phantom. Finite-element studies of helmet therapy have either represented the helmet as boundary constraints and reported intracranial rather than helmet–head contact pressures [10], or modeled a helmet in deformational plagiocephaly under head weight alone [11]; neither was validated against physical measurements. Experimental head–orthosis pressures have so far been reported only for static, rigid head molds in positional plagiocephaly, with peak holding pressures of 8.5–20.5 kPa [15]. The highest local reading here (269 gf, about 52 kPa at B3) was a normal force at one discrete site rather than the total force exerted by the helmet. Its higher value likely reflects differences between the studies: discrete 8-mm contact elements rather than thin-film maps averaged over neighboring cells, and a post-surgical craniosynostosis helmet on an expandable phantom rather than positional-plagiocephaly orthoses on rigid molds. Forces reported for adult protective helmets [14] allow only an approximate comparison, because their users, materials, loading durations and purposes differ fundamentally from postoperative infant orthoses.

Comparing Phantom 3 with and without the helmet shows how the treatment works through contact and relief regions. Applying the helmet to the post-craniotomy phantom constrained displacement in the frontal, occipital and lateral regions, where the sensors recorded contact, and redirected expansion toward the superior clearance, which was largely filled by 175 days (860 mL). Together with the later development of positive force at initially relieved sensing sites, these observations support the complementary roles of retaining contact and relief space: contact regions constrain expansion, while regions left free provide a path into which expansion can be redirected.

Repeated fitting reproduced the loading pattern but also showed the practical limits of measuring force at a small number of sites inside a clinical helmet. The gradual decrease at C1 and A3 over the series most likely reflects small differences in helmet seating between fittings, together with settling of the foam liner, skin or contact elements under repeated loading, rather than a change in the expansion of the phantom.

The study complements rather than replaces computational modeling: the companion finite-element model [13] supplied only the 77-day geometry of Phantoms 2 and 3, and the present experiments were not designed as a point-by-point validation of a particular computational pressure field. Their value is to establish a physical force scale, a spatially non-uniform loading pattern and a directly observed sequence from clearance to contact against which future models can be compared. The setup also provides an experimental route for testing future helmets, material and surgical-design assumptions.

Several limitations should be considered. The study represents one infant, one unicoronal geometry and the first clinical helmet only. Twelve discrete sites cannot provide a continuous pressure map, and the sensors primarily quantify local normal force; shear loading and off-axis sensitivity were not fully characterized. Although Phantoms 2 and 3 shared the same 77-day source geometry and differed only at the fused or released coronal suture, they were separate physical builds and may retain some manufacturing variability. The phantom reproduces controlled elastic expansion rather than biological growth, remodeling, healing or tissue adaptation, and the validation concerns the external skin surface. Force and CT data came from corresponding but separate experiments, so force and displacement were not compared sensor by sensor. There was no *in vivo* force ground truth, and residual uncertainty remains from calibration, manual tare and fitting-dependent sensor seating. Target volumes at the clinical time points were derived from a patient-anchored normative growth curve rather than measured, and the 175-day (860-mL) step represents an extended end of Helmet 1 rather than a recorded clinical time point.

Future work should extend the approach to denser sensing arrays, combined normal and shear-force measurement, calibration under controlled off-axis loading, and registered acquisition of force and three-dimensional cranial shape. Replication across additional patients, synostosis patterns and helmet geometries will be necessary to establish how generalizable the observed local-force scale, regional loading patterns and sequence from clearance to contact are. Patient-derived *in vitro* testing could additionally be used to compare candidate surgical-release geometries, liner-clearance strategies and sequential helmet designs under controlled conditions.

Translated to *in vivo* use, the approach could support longitudinal monitoring of helmet fit, flagging loss of intended contact, premature engagement of relief regions or unexpected focal loading between appointments [17,23,24]. Clinical use would, however, require validated relationships between measured force, skin tolerance and morphological response, and hardware that tolerates months of wear, cleaning, perspiration and hair at the interface [7,8]. Because hair between the liner and sensing surface could alter local load transfer, translation would also require small, low-power and preferably wireless sensors, durable enclosures, preservation of liner compliance and fitting procedures that account for hair, skin condition and repeated helmet placement. *In vivo* sensing should initially complement, not replace, established clinical assessment.

## 4 Conclusion

This study shows that a CT-reproduced clinical helmet can be instrumented to measure local orthotic forces directly, demonstrates that the sensor-level pattern is reproduced across repeated helmet fittings, and shows how the readings vary between fittings when a small number of sensors measure a complex contact surface. By combining direct force measurement with CT-derived shape and clearance observations, the approach shows step by step how contact regions and planned relief space redirect expansion in a patient-derived cranial model. The same approach could be used to test new helmet designs and, in the future, to support clinical interventions, including surgical planning, on patient-derived models.

## 5 Experimental Section

### 5.1 Clinical Data and Ethics

This study was an *in vitro* investigation with repeated measurements, based on clinical data from a single female infant diagnosed with left unicoronal craniosynostosis. The clinical timeline is summarized in Figure 2 and in Section S1, Table S1 (Supporting Information). A CT scan including cranial and external soft-tissue geometries was acquired at 56 days of age, and a three-dimensional surface scan was acquired at 76 days, one day before surgery. To anonymize the clinical data before use, identifying facial features were removed from the external surface meshes: the eye and nasal regions were smoothed in Meshmixer and the projecting ears were cut away, while the helmet-contact region and the upper ear attachment points used for helmet placement were retained. All experiments were performed on physical phantoms, and no experimental intervention was conducted on the infant. Use of the anonymized CT data was approved by University College London (project no. 19055.001). Written informed consent for research use and publication of the imaging data was obtained from the parent(s) or legal guardian(s).

### 5.2 Phantom Configurations and Target Volumes

Phantom 1, made from the patient’s 56-day CT, was used only for longitudinal validation (Sections S5 and S8, Supporting Information). Phantoms 2 and 3, used in the main study, represented the same 77-day (704-mL) surgery-day baseline and were manufactured from full-scale ICV, bone, suture and skin geometries derived from the subject-specific finite-element growth model developed in a companion study, in which the patient’s 56-day geometry was grown to the 77-day surgery-day state [13,25]. Phantom 2 (no operation) retained the fused left coronal suture: the left frontal and parietal bones, including the intervening fused region, formed a continuous rigid component. Phantom 3 (post-operation) was generated from the same 77-day geometry after a surgeon-defined craniotomy had been created in Avizo 2022.1 (Thermo Fisher Scientific, USA) by removing a defined segment of bone across this frontal-parietal suture and was used both with and without the helmet. Intracranial volume was the controlled experimental variable, and each clinical time point was translated into a target volume. Target volumes were derived from a normative intracranial-volume–age relation for females: a cubic polynomial fitted to the female 0–12-month subset of normative data sourced from Liang et al. [26] and Samiepour et al. [13] and reanalyzed here (V = 434 + 117t ™ 11.7t^2^ + 0.527t^3^, with V in mL and t in months; Section S2, Figure S2a, Supporting Information), shifted vertically by a constant offset of +40.4 mL so that it passed through the 704-mL surgery-day baseline at 77 days, following the approach of Samiepour et al. [13]. The full equation and conversion table are provided in Section S2, Table S2 (Supporting Information). This gave target volumes at surgery, 77 days (704 mL); at the clinical fitting of Helmet 1, 90 days (732 mL); and at its end, 160 days (843 mL). Because the normative volumes are population references and the end of a helmet period is adjusted clinically, the end of Helmet 1 was represented by a range rather than a single value: in addition to 160 days (843 mL), a step at 175 days (860 mL) was included and imaged by CT.

At this volume, the helmet clearance was visually filled (Figure 7). A further extension to 212 days (900 mL), beyond the Helmet 1 period, tested the response when expansion continued and is reported in Sections S7 and S8 (Supporting Information). Helmet 1 was manufactured from the patient’s 76-day pre-operative surface scan and, because clinical fabrication takes one to two weeks, was first worn at 90 days. *In vitro*, the instrumented helmet was fitted from the surgery-day baseline (77 days, 704 mL) so that the onset of contact, including any contact before the clinical fitting, was captured. The phantoms, helmets and sensor sets used in each experiment are listed in Section S5, Table S3 (Supporting Information).

### 5.3 Phantom Fabrication and Helmet Instrumentation

All phantoms were assembled using the sacrificial-core, multi-material process established in the preceding *in vitro* platform [22]. In brief, CT- or model-derived intracranial-volume cores, cranial bone plates individually manufactured using stereolithography (SLA) printers, compliant latex sutures and inner and outer latex walls were combined to form hollow expandable skull constructs. For Phantom 2, the fused left frontal-parietal unit was produced as one continuous stereolithography-printed rigid component, so the fused suture could not open during expansion. For Phantom 3, no rigid material was printed within the surgeon-defined resection region. During assembly, this opening was filled with the same compliant latex used for the sutures and was integrated with the surrounding inner and outer latex layers, preserving structural continuity while allowing the modeled post-craniotomy region to expand. A patient-conformal Ecoflex 00-50 silicone skin (Smooth-On, USA) was molded over each completed skull. The phantoms were connected through the foramen magnum (the opening at the skull base) to the previously described pneumatic–hydraulic expansion system. Full material preparation, assembly, layer-formation, skin-molding and pre-use integrity procedures are provided in Sections S3 and S4, Figure S3 (Supporting Information).

The full-scale geometry of the patient’s first clinical orthosis (Helmet 1) was acquired by computed tomography using an XT H 225 ST system (Nikon Metrology, Tring, UK) at 150 kV, 150 µA, with a voxel size of 50 µm and 2,000 projections. The shell was segmented in Avizo 2022.1 (Thermo Fisher Scientific, USA) and modified for sensor integration using Fusion 360 and Meshmixer (Autodesk, USA) and SOLIDWORKS (Dassault Systèmes, France). Openings and fixation features were added while preserving the 1:1 clinical scale, and the modified outer shell was printed at full scale in acrylonitrile butadiene styrene (ABS) on an H2D printer (Bambu Lab, China) using the manufacturer’s standard ABS profile. A new 3-mm Plastazote LD33 liner (Zotefoams, UK; supplied by Algeos, UK) was selected to match the material and nominal thickness of the clinical helmet liner and was bonded to the inner surface of the printed shell. The liner was a closed-cell, cross-linked polyethylene foam with a nominal density of 33 kg m™3. The resulting helmet is a full-scale (1:1) copy of Helmet 1 with the same size and shape, apart from the added sensor openings and fixation features. Fitted with the twelve force sensors described below, it is referred to throughout as the instrumented helmet (Figure 3b, d).

Twelve custom soft magnetic tactile force sensors based on Hall-effect sensing were integrated at clinically selected locations determined with input from the treating surgeon and assigned a priori to three anatomical groups: frontal (A1–A4), occipital (B1–B4) and bilateral spheno-temporal (C1–C2 left, C3–C4 right). The sensor design was adapted from previously reported soft tactile-sensor architectures [17,27,28]. Each sensor comprised a Hall-effect printed circuit board, a neodymium magnet, an 8-mm-diameter and 3-mm-high cylindrical elastomeric contact element, and a stereolithography-printed mount. Sensors were mounted through the ABS shell between the shell and foam liner without protruding beyond the inner foam surface, providing 12 discrete local force-measurement sites. Three four-channel bus hubs connected the array to a Teensy 4.1 microcontroller (PJRC, USA), and Sensor Tracking Studio acquired and displayed the 12 channels in real time. Each sensor was calibrated individually against an ATI Nano17 six-axis force/torque transducer (ATI Industrial Automation, USA) on a custom three-axis platform using motorized translation stages (Thorlabs, USA; Section S9, Video S1, Supporting Information), and a sensor-specific third-order (cubic) force-conversion equation was fitted to the loading branch of the third of five calibration cycles. A cubic was chosen because a second-order fit over-predicted low forces and produced spurious readings at non-contacting sites. The complete sensor-fabrication, calibration, signal-processing and calibration-performance analyses are provided in Sections S6.1–S6.5 (Supporting Information).

Before each run, sensor-to-channel assignment was verified by light manual pressure on each sensor (Section S9, Video S2, Supporting Information). Sensors were not removed, repositioned or recalibrated during the series.

### 5.4 Experimental Protocol

The main study comprised three configurations evaluated from 77 days (704 mL) to 175 days (860 mL): Phantom 2 without a helmet, Phantom 3 without a helmet and Phantom 3 with the helmet. Comparing Phantom 3 with and without the helmet isolated the effect of the helmet on shape, whereas Phantom 2 showed the effect of the modeled craniotomy.

Before each force-measurement run, the phantom and hydraulic line were filled with water, evacuated to minimize trapped air and prepared at the 77-day (704-mL) baseline. For force experiments, the sensors were manually tared in the unloaded state before the instrumented helmet was fitted to the phantom. Water was delivered using the previously described pneumatic–hydraulic expansion system; when a prescribed volume was reached, the hydraulic valve closed automatically, and the system was held at that volume until the corresponding recording phase was complete [22]. For CT imaging, the corresponding mechanical configurations were reproduced separately. The pneumatic–hydraulic accumulator was not used in the scanner; iodinated contrast solution was introduced and withdrawn through the inlet at the foramen magnum of the phantom base using graduated 100- and 200-mL syringes to remove trapped air and establish the prescribed volumes. In the helmeted CT condition, a dummy helmet was used, a full-scale, metal-free copy of the instrumented helmet with identical geometry that contained no magnets, printed circuit boards, electronics or other metallic components; dimensionally matched non-sensing inserts and plastic fasteners replaced the active sensors and original metal fasteners. Dummy-helmet placement used the same landmarks and strap setting as the force experiments. Because of the substituted inserts and fasteners, exact equivalence of local seating was not assumed; placement was accepted when the overall helmet–skin contact pattern and designed relief gaps showed no visible discrepancy from the instrumented configuration.

Helmet placement was standardized using three anatomical landmarks on the skin geometry: an anterior midline reference at the glabella (the point between the eyebrows) and the upper attachment points of both ears (otobasion superius). Strap fastening was standardized using a visual reference mark on the strap to reproduce the same nominal strap setting at every fitting.

Before fitting, a light dusting of talc powder was applied at the interface between the silicone skin and the elastomeric sensor contact elements to reduce adhesion between the two elastomeric surfaces. This limited sticking, folding and lateral drag of the contact elements during fitting. Force and raw Bz channels were monitored live in Sensor Tracking Studio. If the baseline pattern indicated a gross seating inconsistency, visible off-axis deformation of a contact element or an implausible sensor response, the run was stopped and the helmet was removed and refitted. The experiment was then repeated from the 77-day (704-mL) baseline.

Force experiments with the instrumented helmet comprised seven expansion runs. Consecutive runs were separated by approximately 1–2 min to release the volume and to remove and refit the helmet; the series also contained two longer unloaded intervals of approximately 1 h and 1.5 h, introduced to allow the elastomeric contact elements to recover, so that fittings performed in immediate succession and after recovery were both represented. The helmet was removed and refitted before every run, so that each run represented an independent fitting. The seven runs (n = 7) form the dataset reported here, and repeatability describes how closely the sensor-level force pattern was reproduced across these repeated fittings. The preparation and expansion run of Fitting 6 are shown in Section S9, Videos S2 and S3 (Supporting Information).

### 5.5 CT Imaging and Surface Analysis

Phantoms were imaged at selected target volumes using an XT H 225 ST system (Nikon Metrology, Tring, UK), filled with the contrast solution described above; water was used in all other experiments. Reconstruction, segmentation and extraction of external surface meshes were performed in Avizo 2022.1 (Thermo Fisher Scientific, USA).

For baseline validation, the external skin surfaces of Phantoms 2 and 3 at 77 days (704 mL) were compared with the patient’s 76-day pre-operative surface scan; the longitudinal validation with Phantom 1 is described in Section S8, Figure S14 (Supporting Information).

Clinical and phantom surface meshes were registered in a common coordinate system using the automatic alignment tool in Avizo. For validation, surface differences were assessed using surface-distance maps with a common 0–5-mm color scale. For each mechanical configuration, CT surfaces acquired at 77 and 175 days (704 and 860 mL) were compared to visualize three-dimensional shape change over the main experimental interval. Helmet–skin clearance and its reduction during intracranial-volume expansion were also assessed in four standard section planes: P1 sagittal, P2 and P3 coronal, and P4 horizontal, using overlaid helmet, 77-day (704-mL) and 175-day (860-mL) skin geometries.

### 5.6 Force Measurement and Data Processing

Force measurement was based on the load-dependent change in distance between a magnet embedded in the elastomeric contact element and the Hall-effect sensor [27]. Compression of the elastomer changed the raw magnetic-field output, which was converted to force with the sensor-specific cubic equation described above. Calibration was performed under loading normal to the sensor face; sensitivity to off-axis magnet displacement was not independently characterized.

Forces are reported in gram-force (gf; 1 gf = 9.807 mN); values in newtons and nominal pressure at 175 days (860 mL) are given in Section S7, Table S6 (Supporting Information). Nominal sensor-face pressure was calculated, where reported, by dividing local force by the nominal circular face area of the 8-mm-diameter contact element (50.3 mm^2^). The conversion assumes uniform full-face contact and does not represent a spatially resolved interface-pressure measurement.

During force experiments, volume was increased stepwise through the target volumes and then returned to the baseline volume (704 mL), with each target volume held while it was recorded. The operator labeled each acquisition phase in Sensor Tracking Studio at its onset, distinguishing stable target volumes (for example, the 90-day (732-mL) hold) from transitions (for example, from 704 to 732 mL), and the active label was written to every timestamped data row. For each run and sensor, the force at a target volume was calculated as the mean of all samples in the first stable segment carrying that volume label, so that the 77-day (704-mL) value always refers to the pre-loading baseline.

For regional analysis, per-sensor normal-force magnitudes at a given volume were summed within each predefined anatomical group and expressed as a percentage of the summed magnitudes across all twelve sensing sites in the same run. These quantities represent sampled regional load shares and not vector-resultant forces or the total reaction force of the helmet.

### 5.7 Statistical Analysis

No data were removed as outliers and no normalization was applied. Data are presented as mean ± standard deviation (sample SD) across the seven fittings (n = 7). Fitting repeatability was described by the standard deviation and coefficient of variation of each sensor’s force across fittings and by the range of regional load shares; sensor-level repeatability was assessed on the calibration rig by repeated normal loading and is reported in Section S6.4, Figure S11 and Table S4 (Supporting Information). All analyses were descriptive, and no inferential statistical tests were applied. Data processing and plotting were performed in Python 3 (pandas, NumPy and Matplotlib).

## Supporting Information

Supporting Information is available from the Wiley Online Library or from the author.

## Acknowledgements

We thank W. Stokes, whose needle-insertion rig software was the starting point for the calibration-rig software. Ahmet Arif Colakoglu is supported by a scholarship from the Ministry of National Education of the Republic of Türkiye. Mehran Moazen is supported by the Engineering and Physical Sciences Research Council (EP/W008092/1). During the preparation of this work, the authors used Claude (Anthropic) and ChatGPT (OpenAI) to assist with language editing. The authors reviewed and edited the AI-assisted text and take full responsibility for the content of the publication.

## Author Contributions

All authors contributed to the conception and design of the study. Material preparation, experimental setup development, experiments and analysis were performed by Ahmet Arif Colakoglu under the supervision of Peter Culmer and Ali Alazmani. The first draft of the manuscript was written by Ahmet Arif Colakoglu, and all authors commented on previous versions of the manuscript. All authors read and approved the final manuscript.

## Data Availability Statement

The force logs of the seven fittings, the calibration step medians of the twelve sensors and the scripts that reproduce Figure 5 and, in the Supporting Information, Figures S9–S12 and Table S4 are available at https://github.com/aarifcolakoglu-research/helmet-contact-force-analysis (archived at https://doi.org/10.5281/zenodo.24340880). The source code of Sensor Tracking Studio, including the calibration pipeline, the equation set used in this study and the Teensy 4.1 firmware, is available at https://github.com/aarifcolakoglu-research/sensor-tracking-studio (https://doi.org/10.5281/zenodo.24340917), and the calibration-rig software, version 1.0 of which recorded the sensor calibrations, at https://github.com/aarifcolakoglu-research/tactile-sensor-calibration-rig (https://doi.org/10.5281/zenodo.24340958). The raw calibration recordings (about 2.2 GB) and other data generated and analyzed during the current study are available from the corresponding author upon reasonable request.

