## Supporting Information for "Evaluation of an Instrumented Helmet for Measurement of Orthosis–Head Interface Forces in Pediatric Craniosynostosis"

#### S1. Clinical Data and Treatment Time Line

The clinical data come from one female infant with left unicoronal craniosynostosis who had surgery at 77 days of age and then wore three cranial orthoses in turn (Figure 2, main text; Table S1). Figure S1 shows the clinical helmets and the replicas of Helmet 1 made for this study. Helmet 1 was made from the 76-day surface scan and first worn at 90 days; *in vitro*, the instrumented helmet was fitted from surgery (77 days) to capture the onset of contact. Only the first treatment phase, from surgery to the end of Helmet 1, was represented *in vitro*, with an extended end-of-Helmet-1 step at 175 days (860 mL), at which *in vitro* CT was acquired, and an extension to 212 days (900 mL) reported only in this Supporting Information; later phases and follow-up are listed for context.

**Table S1.** Complete clinical time line of the patient. Rows shaded in amber fall within the interval represented *in vitro* (77–175 days).

| Age [days] | Clinical data | Treatment / evaluation | Helmet worn | Head circumference [cm] |
|---|---|---|---|---|
| 56 | CT (skull and skin) | – | – | – |
| 76 | 3D surface scan (skin) | Pre-operative evaluation | – | 38.9 |
| 77 | – | Surgery; instrumented helmet fitted *in vitro* from this point | – | – |
| 90 | – | Helmet 1 fitted clinically (worn 90–160 days, 70 days; represented *in vitro*) | Helmet 1 | – |
| 160 | – | End of Helmet 1; Helmet 2 fitted (worn 160–320 days, 160 days; outside the scope) | Helmet 1 → 2 | – |
| 320 | – | End of Helmet 2; Helmet 3 fitted (worn 320–377 days, 57 days; outside the scope) | Helmet 2 → 3 | – |
| 377 | 3D surface scan (skin) | Follow-up evaluation; end of Helmet 3 | Helmet 3 | 44.6 |
| 482 | 3D surface scan (skin) | Follow-up evaluation | – | 45.7 |
| 566 | 3D surface scan (skin) | Follow-up evaluation | – | 46.5 |
| 727 | 3D surface scan (skin) | Follow-up evaluation | – | 46.8 |
| 1518 | 3D surface scan (skin) | Follow-up evaluation | – | 49.0 |
| 1729 | 3D surface scan (skin) | Follow-up evaluation | – | 49.2 |
| 3456 | 3D surface scan (skin) | – | – | – |
| 3730 | – | Follow-up evaluation | – | 51.0 |

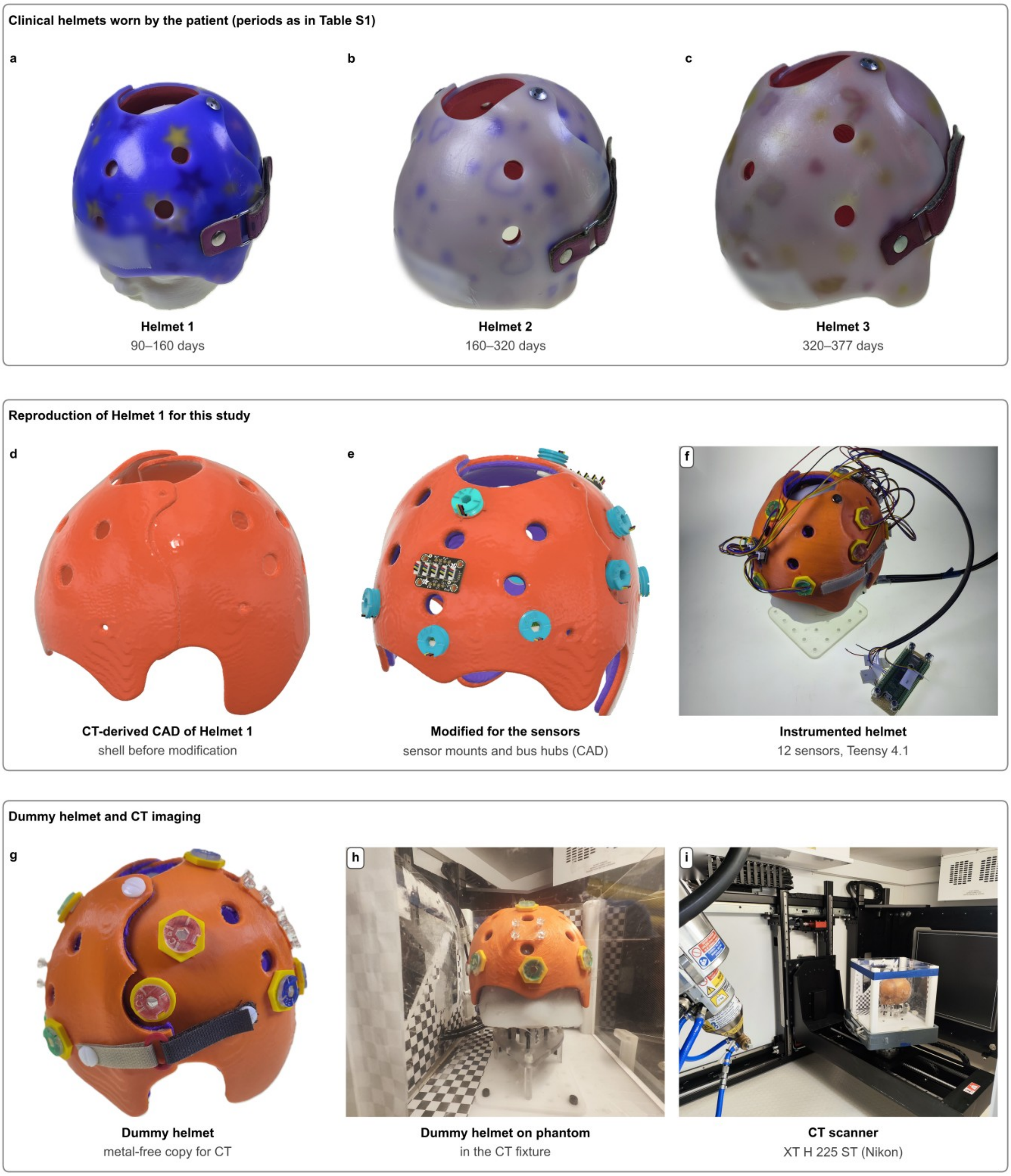


**Figure S1.** Clinical helmets and replicas. (a–c) Helmets 1–3 worn by the patient. (d) CT-derived CAD model of Helmet 1. (e) Model modified for sensor mounting. (f) Instrumented helmet on the phantom. (g) Dummy helmet. (h) Dummy helmet on the phantom in the CT fixture. (i) CT scanner. Handwritten labels are blurred.

CT imaging covered 77, 175 and 212 days (704, 860 and 900 mL) but not 90 and 160 days (732 and 843 mL). The CT volumes were selected before the clinical target volumes had been derived from the normative relation (Section S2): 860 mL was chosen as the volume at which the phantom visibly filled Helmet 1, and 900 mL was added as an extension. Under the shifted relation, 860 mL corresponds to 175 days; because the normative volumes are population references and the end of a helmet period is adjusted clinically, the main study treats 175 days (860 mL) as an extended end-of-Helmet-1 step, shortly after the clinical end of Helmet 1 at 160 days (843 mL). The force experiments covered all five volumes.

#### S2. Target Intracranial Volumes

Intracranial volume (ICV) was the controlled experimental variable. Target volumes at the clinical time points were derived from a normative ICV–age relation for females: a cubic polynomial fitted to the female 0–12-month subset of normative data sourced from Liang et al. [1] and Samiepour et al. [2] and reanalyzed here:

$$V(t) = 434 + 117t - 11.7t^2 + 0.527t^3,$$

where $V$ is the ICV in mL and $t$ is age in months, with ages in days converted at 30.4375 days per month (365.25/12). The female fit has $R^2 = 0.86$ ($P < 0.001$; Figure S2a); the corresponding male fit is shown for comparison only and was not used. The unshifted relation gives 663.6 mL at 77 days. Following the constant-offset approach of Samiepour et al. [2], the relation was shifted vertically by +40.4 mL so that it passed through the 704-mL surgery-day baseline of Phantoms 2 and 3 at 77 days; the 704-mL baseline is the ICV of the 77-day geometry supplied by the companion finite-element model [2,3]. The shifted relation was then evaluated at the clinical time points and rounded to the nearest milliliter (Table S2, Figure S2b). The 56-day, 572-mL starting volume of Phantom 1 was measured directly from the segmented patient CT and was not derived from the shifted relation. The relation was fitted to 0–12-month data and was not used beyond this range.

**Table S2.** Conversion of clinical ages to target intracranial volumes.

| Age [days] | Age [months] | Unshifted relation [mL] | Shifted relation (+40.4 mL) [mL] | Target volume used [mL] | Role |
|---|---|---|---|---|---|
| 56 | 1.84 | 612.9 | 653.3 | 572 | Phantom 1 baseline (measured from CT) |
| 77 | 2.53 | 663.6 | 704.0 | 704 | Surgery; baseline of Phantoms 2 and 3 |
| 90 | 2.96 | 691.3 | 731.7 | 732 | Clinical fitting of Helmet 1 |
| 160 | 5.26 | 802.3 | 842.7 | 843 | End of Helmet 1 |
| 175 | 5.75 | 820.1 | 860.5 | 860 | Extended end of Helmet 1 (CT) |
| 212 | 6.97 | 859.4 | 899.8 | 900 | Extension (Supporting Information only) |

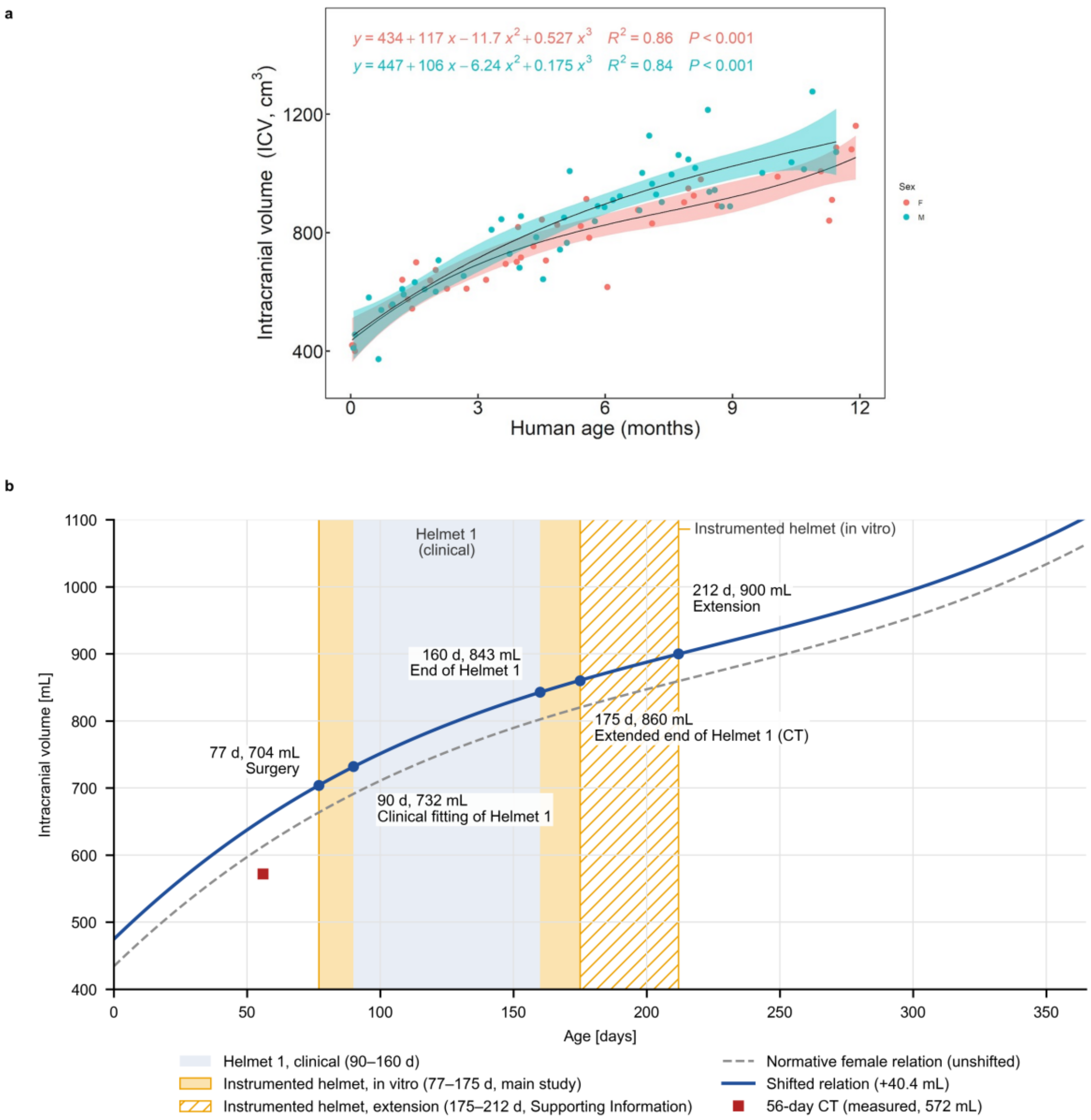


**Figure S2.** Target intracranial volumes. (a) Third-order fits to the 0–12-month normative ICV data sourced from Liang et al. [1] and Samiepour et al. [2] and reanalyzed here (female, red; male, teal; shaded, confidence bands). (b) Female relation before (dashed) and after (solid) the +40.4-mL shift; circles, target volumes; square, 572 mL measured from the 56-day CT; blue band, clinical Helmet 1 (90–160 days); amber, instrumented helmet *in vitro* (77–175 days; hatched, 212-day extension).

### S3. Phantom Fabrication and Integrity Testing

All three phantoms were made with the method of our previous *in vitro* platform, which uses a dissolvable core and layers of latex [4]. For each phantom, the patient-derived ICV geometry was printed in water-soluble polyvinyl alcohol (PVA) on an X1-Carbon printer (Bambu Lab, China) to form a temporary assembly core, and the individual cranial bone plates were manufactured using stereolithography (SLA) printers. The bone plates were placed on the PVA core and held in position with pins inserted through reference holes; for the post-craniotomy phantom, the surgeon-defined craniotomy was added at this stage (Figure 3a, main text; Figure S3h–j, p–r).

The suture gaps were filled by syringe with liquid latex (Polycraft Liquid Latex – Brush On/Dipping Latex, MB Fibreglass, UK) containing 5% by mass of latex thickener (Polycraft Latex Thickener, MB Fibreglass, UK); the components were weighed using a precision balance. The assembly was allowed to air-dry for 12 h. The registration pins were then removed and the complete skull was dip-coated in the same liquid latex without thickener. Successive coats were applied in a ventilated laboratory at 25 °C, each coat being allowed to dry before the next, until at least three external layers had been formed. These layers held the separate bone plates together as one skull. Small holes (2 mm) across the bone plates let the latex pass through the plates and later join the inner and outer latex walls.

The PVA core was dissolved through the foramen magnum (the opening at the skull base) in a water-filled ultrasonic bath maintained at no more than 35 °C to avoid thermal damage to the printed components and latex. This took about 24 h and was continued until no PVA was visible. The hollow skull was then filled with unthickened liquid latex and drained through the same opening; each internal coat was allowed to dry at 25 °C, and the cycle was repeated at least three times. Further dipping and fill-and-drain cycles built up both walls until each surface had nine latex layers, giving a sealed, flexible expansion chamber. Latex tubing was attached at the foramen magnum to connect the phantom to the expansion system. The layer build-up had been checked earlier by cutting a separate test phantom and measuring the latex thickness with calipers; all study phantoms were made with the same coating steps. Before use, each phantom was expanded without a helmet and inspected; it was accepted only if it showed no visible leak or damage.

Expansion in the force experiments and volume setting for CT followed the procedures described in Sections 5.4 and 5.5 (main text). A CT acquisition required approximately 30–35 min, and each target-volume CT stage, including preparation and post-scan checks, approximately 50–55 min. Each force run, from the 77-day (704-mL) baseline through 90, 160, 175 and 212 days (732, 843, 860 and 900 mL) and back to 704 mL, lasted approximately 6.5–9 min.

### S4. Silicone Skin

The skin was cast from Ecoflex 00-50 (Smooth-On, USA), a very soft platinum-cure silicone rubber (Shore hardness 00-50, 100% modulus 0.083 MPa, elongation at break 980% [5]). The skin geometry was the patient’s own external head surface over the corresponding skull; the cranial vault, which carries the helmet, was not changed by anonymization. The skin thickness therefore varied over the head, from approximately 1 to 6 mm. For each phantom, the completed latex-coated skull was wrapped in tape to define the inner mold surface, and a two-part outer mold was designed in Fusion 360 (Autodesk, USA) as the negative of the patient-derived external skin geometry and printed in PLA and ABS on an H2D printer (Bambu Lab, China). A locating fixture attached to a top cap held the skull in the correct position and held the two outer mold halves and the skull together as one closed assembly (Figure S3e, l, m). The two silicone components were mixed 1:1 by weight on a precision balance, poured into the cavity between the latex-coated skull and the outer mold, degassed under vacuum for 15 min and left to cure overnight at room temperature (much longer than the manufacturer’s 3-h cure time) before it was removed from the mold.

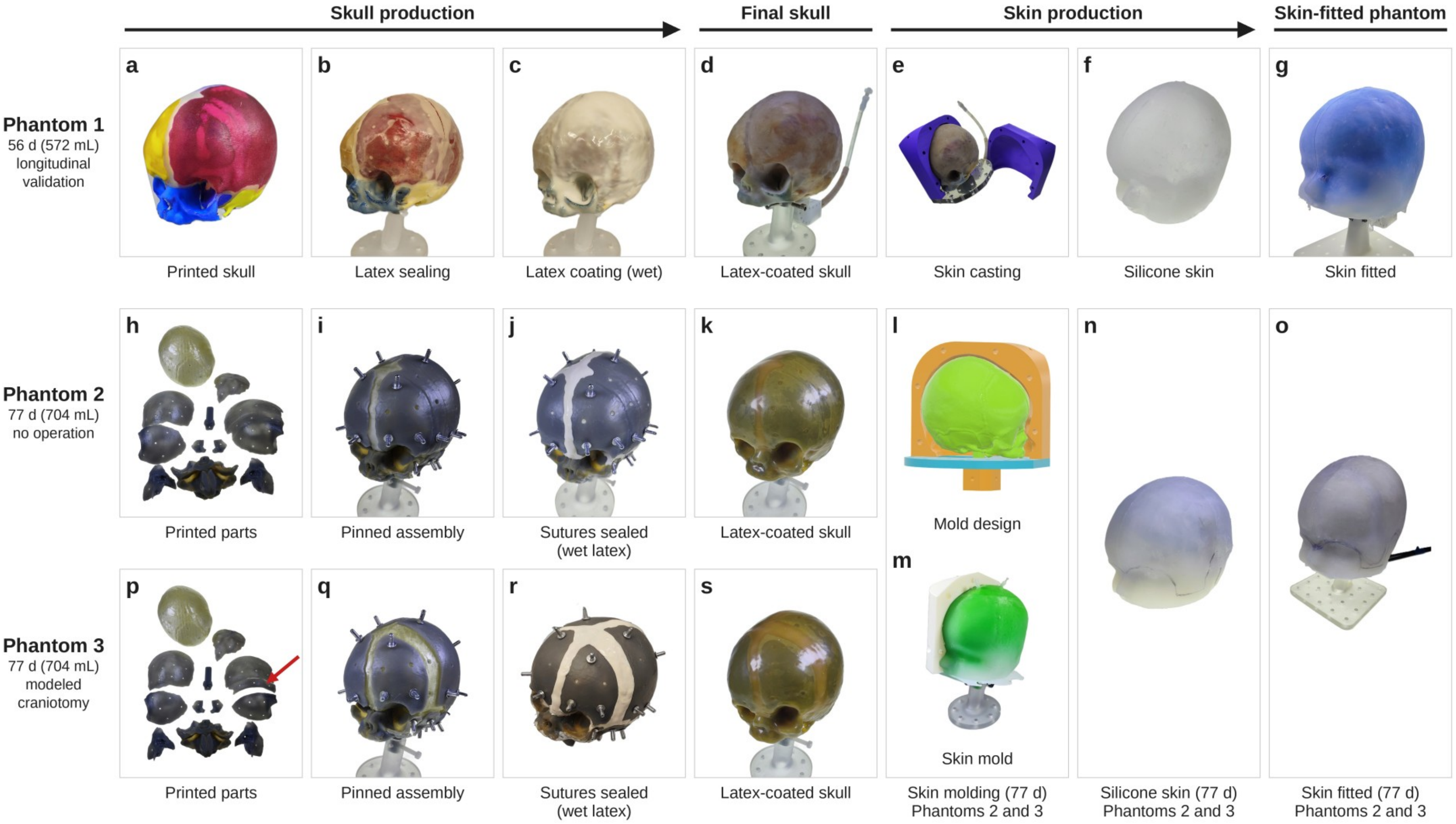


**Figure S3.** Phantom fabrication. Skull production (a–c, h–j, p–r), final latex-coated skulls (d, k, s), skin production (e, f, l–n) and skin-fitted phantoms (g, o) for Phantom 1 (56 days, 572 mL), Phantom 2 (77 days, 704 mL; no operation) and Phantom 3 (77 days, 704 mL; modeled craniotomy, cut marked by the arrow in p). Phantoms 2 and 3 share one skin. Photographs are not to scale.

### S5. Phantoms, Helmets and Experiments

**Table S3.** Phantoms, helmets and sensor set used in each experiment. Photographs of the three phantoms are shown in Figure S3.

| Item | Source geometry and description | Baseline volume | Experiments | Reported in |
|---|---|---|---|---|
| **Phantom 1** | 56-day patient CT; native bone plates, sutures and skin | 56 d (572 mL) | CT at 56 and 77 days (572 and 704 mL); longitudinal validation | Figure S14 |
| **Phantom 2** | 77-day FE-derived geometry; no craniotomy (fused left frontal–parietal unit printed as one rigid component) | 77 d (704 mL) | CT at 77, 175 and 212 days (704, 860 and 900 mL) without helmet; baseline validation | Figures 4, 6; Section S8 |
| **Phantom 3** | 77-day FE-derived geometry; modeled post-craniotomy (resection region bridged with latex) | 77 d (704 mL) | CT at 77, 175 and 212 days (704, 860 and 900 mL) without helmet and with the dummy helmet; baseline validation; seven force runs with the instrumented helmet (77 → 90 → 160 → 175 → 212 days, then release; 704 → 732 → 843 → 860 → 900 → 704 mL) | Figures 4–7; Sections S7, S8 |
| **Instrumented helmet** | 1:1 copy of Helmet 1: ABS shell (H2D, Bambu Lab), 3-mm Plastazote LD33 liner, sensor openings and fixation features; twelve sensors (A1–A4, B1–B4, C1–C4), three bus hubs, Teensy 4.1 | – | All force runs | Figures 3, 5 |
| **Dummy helmet** | Metal-free copy of the instrumented helmet: identical shell and liner geometry, non-sensing inserts and plastic fasteners | – | CT of Phantom 3 with helmet | Figures 6, 7 |
| **Sensor set** | Calibration set S1_MatC: twelve sensors, each with its own cubic calibration; not removed, repositioned or recalibrated during the series | – | All force runs | Section S6 |

### S6. Sensors, Sensor Tracking Studio and Calibration

#### S6.1 Sensor Design and Electronics

The sensor design (Section 5.3, main text) was based on earlier soft tactile sensors [6–8]; the magnet, about 0.8 mm thick, was embedded in the elastomeric contact element. Compression of the elastomer moves the magnet toward the Hall-effect sensor and changes the measured magnetic flux density ($B_z$), which is converted to force. The twelve sensors were divided across three four-channel bus hubs and connected to a Teensy 4.1 microcontroller (PJRC, USA) through an $I^2C$ bus (Figure S4). The microcontroller reports each reading with its socket (A1–C4); which calibrated sensor occupies each socket is defined in Sensor Tracking Studio.

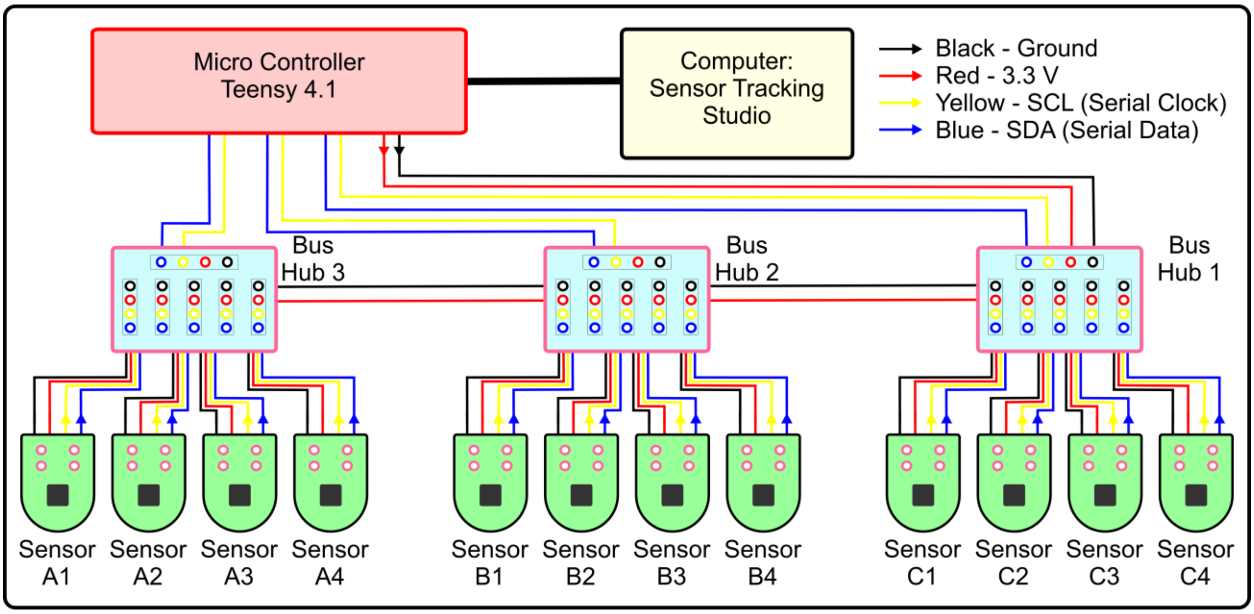


**Figure S4.** Wiring schematic of the 12-sensor array. The sensors are connected in groups of four to three bus hubs, which share the $I^2C$ clock (SCL) and data (SDA) lines, 3.3-V supply and ground of the Teensy 4.1 microcontroller; the microcontroller streams the data to Sensor Tracking Studio over USB.

#### S6.2 Sensor Tracking Studio

Sensor Tracking Studio is a custom Python desktop application (PyQt6) developed for this work that combines the calibration of the sensors, the assignment of calibrated sensors to the twelve sockets and live monitoring and recording in one program [9]. It has three pages (Figures S5–S7):

• **Setup** runs the calibration pipeline (Section S6.4), selects the model used for live readings and maps the twelve sockets to calibrated sensors.

• **Dashboard** shows the raw $B_z$ value and force of each socket, with the recording, volume-label and TARE ALL controls.

• **Overlay** plots all twelve channels on one axis.

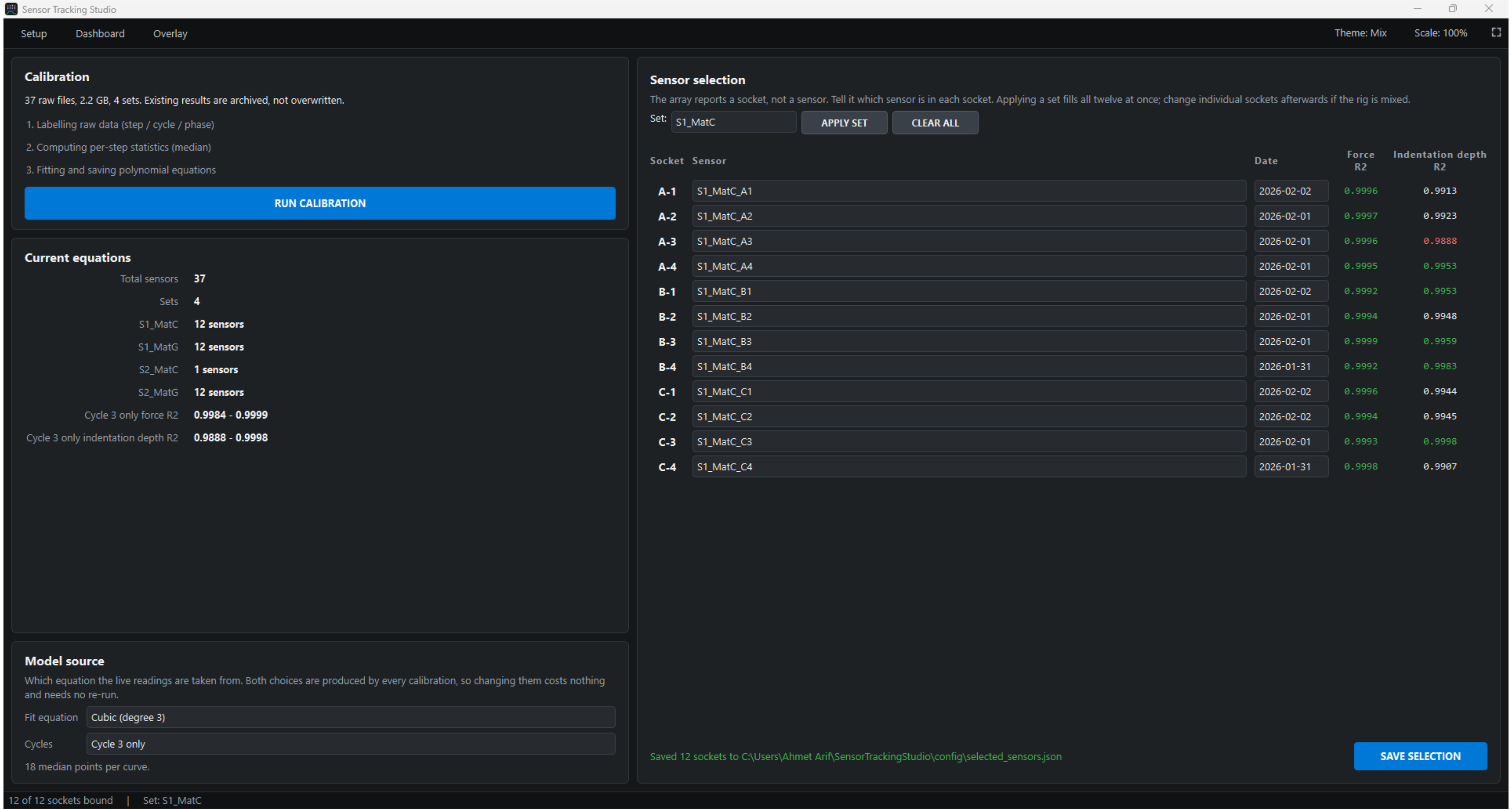


**Figure S5.** Sensor Tracking Studio, Setup page: calibration pipeline, model selection and assignment of the twelve sockets to calibration set S1_MatC.

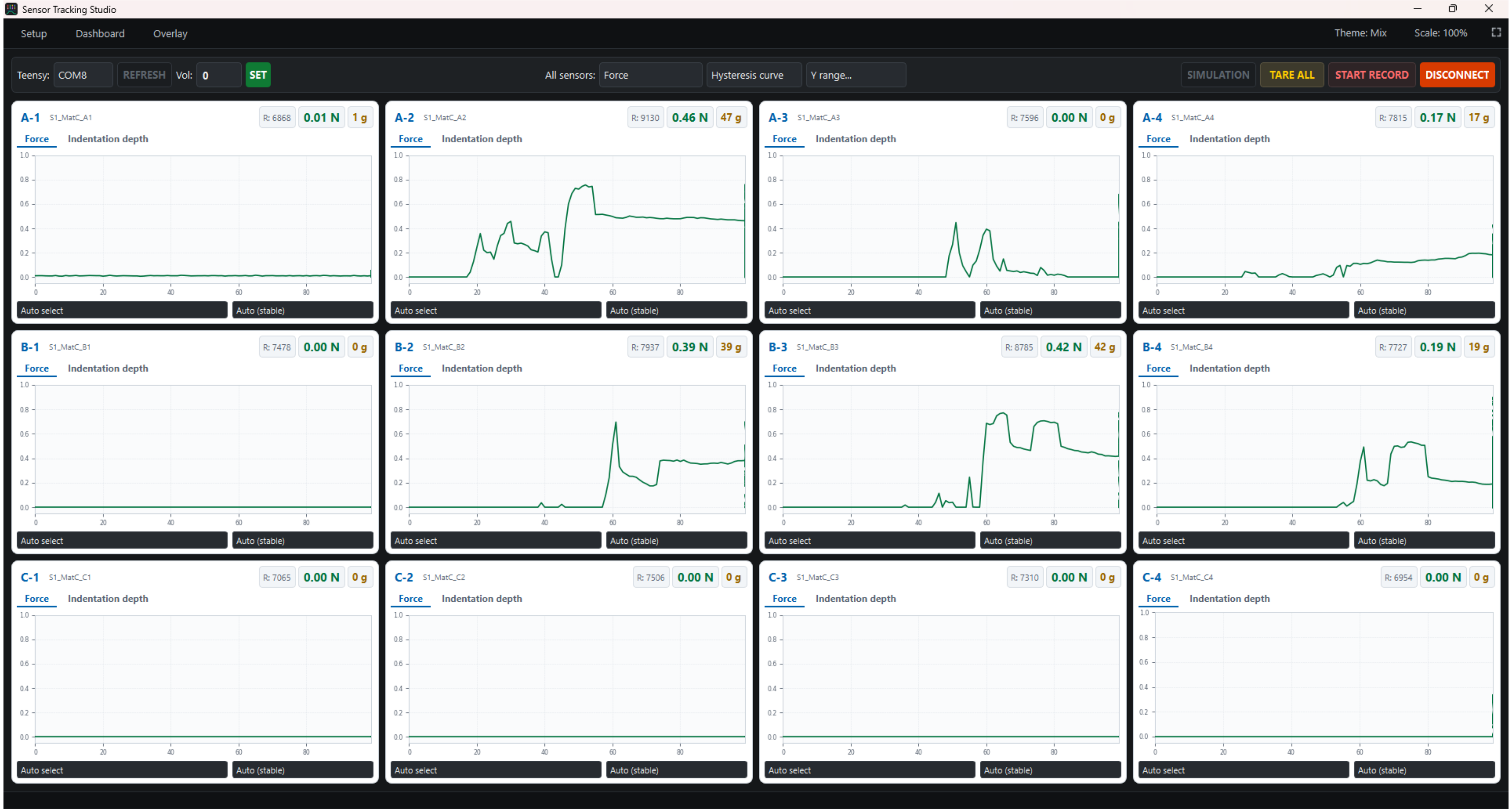


**Figure S6.** Sensor Tracking Studio, Dashboard page during light manual loading of individual sensors.

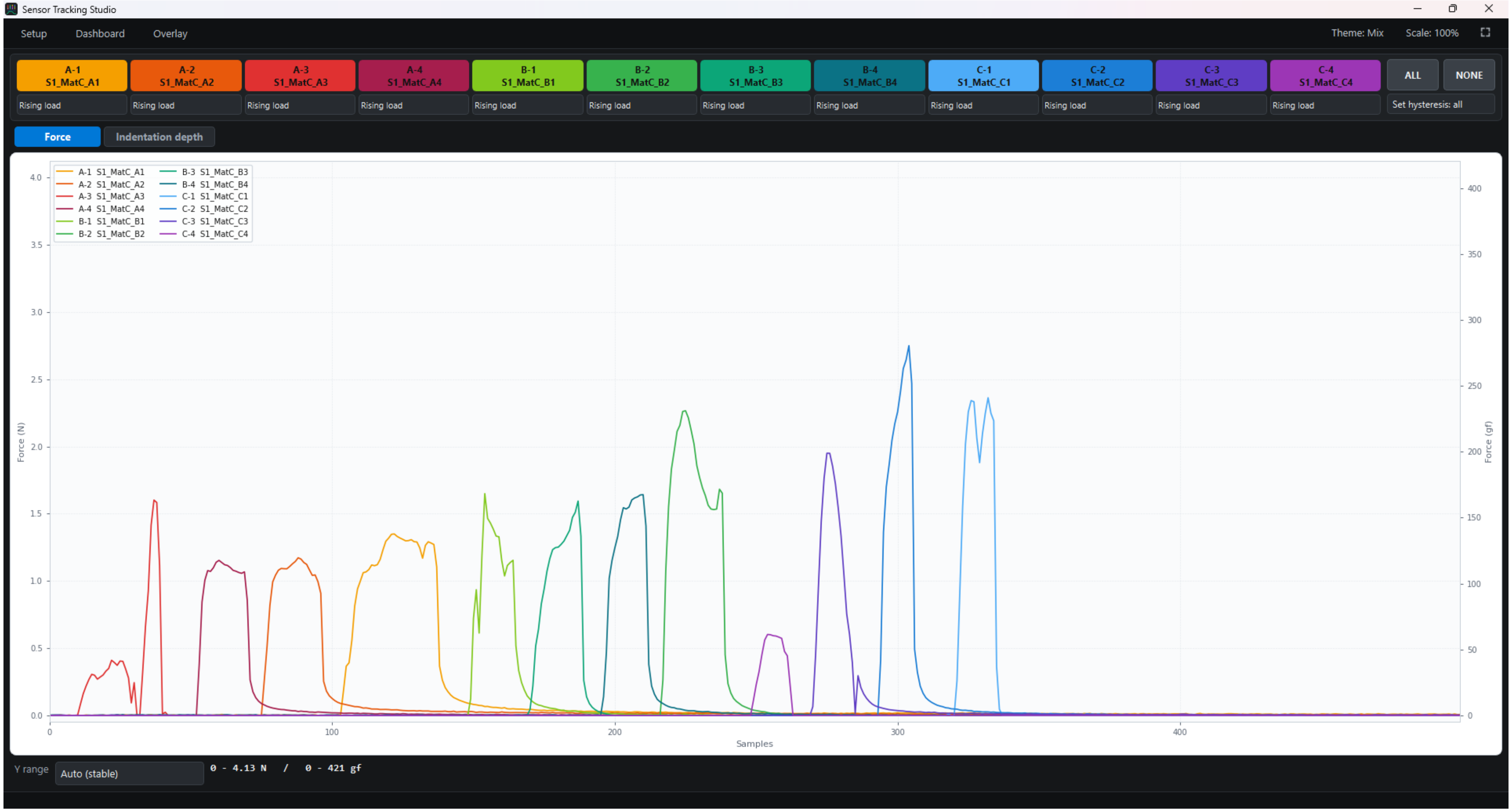


**Figure S7.** Sensor Tracking Studio, Overlay page (force view) while the twelve sensors were pressed in turn.

#### S6.3 Calibration Rig

Each sensor was calibrated individually against an ATI Nano17 six-axis force/torque transducer (ATI Industrial Automation, USA) on a custom three-axis rig built from motorized translation stages (Thorlabs, USA) (Figure S8, Video S1). The sensor was fixed under the transducer in a custom 3D-printed holder, and a probe mounted on the transducer was sized to cover the elastomeric contact element (flat circular tip, 15 mm in diameter; Figure S8b), so that the element was pressed perpendicular to its face. Raw transducer gauge signals and stage encoder positions were acquired by an FRDM-K64F board (NXP Semiconductors, Netherlands), converted to force using the ATI 6 × 6 calibration matrix, timestamped on the device and streamed to the host computer. Raw Hall outputs from the sensor under test were streamed in parallel from a Teensy microcontroller over USB serial at 115 200 baud, and the most recent Hall sample was paired with each transducer record. An automatic contact-detection routine set the zero-indentation position. The $z$ stage then advanced in 0.1-mm increments at 0.5 mm $s^{-1}$ to a maximum indentation of 1.8 mm, with a 10-s hold after every step, before unloading in identical increments. Five loading–unloading cycles were performed per sensor, separated by 20-s rests. Indentation was limited to 1.8 mm because, with a 3-mm elastomer and an embedded magnet of about 0.8 mm, only slightly more than 2 mm of the element can be compressed.

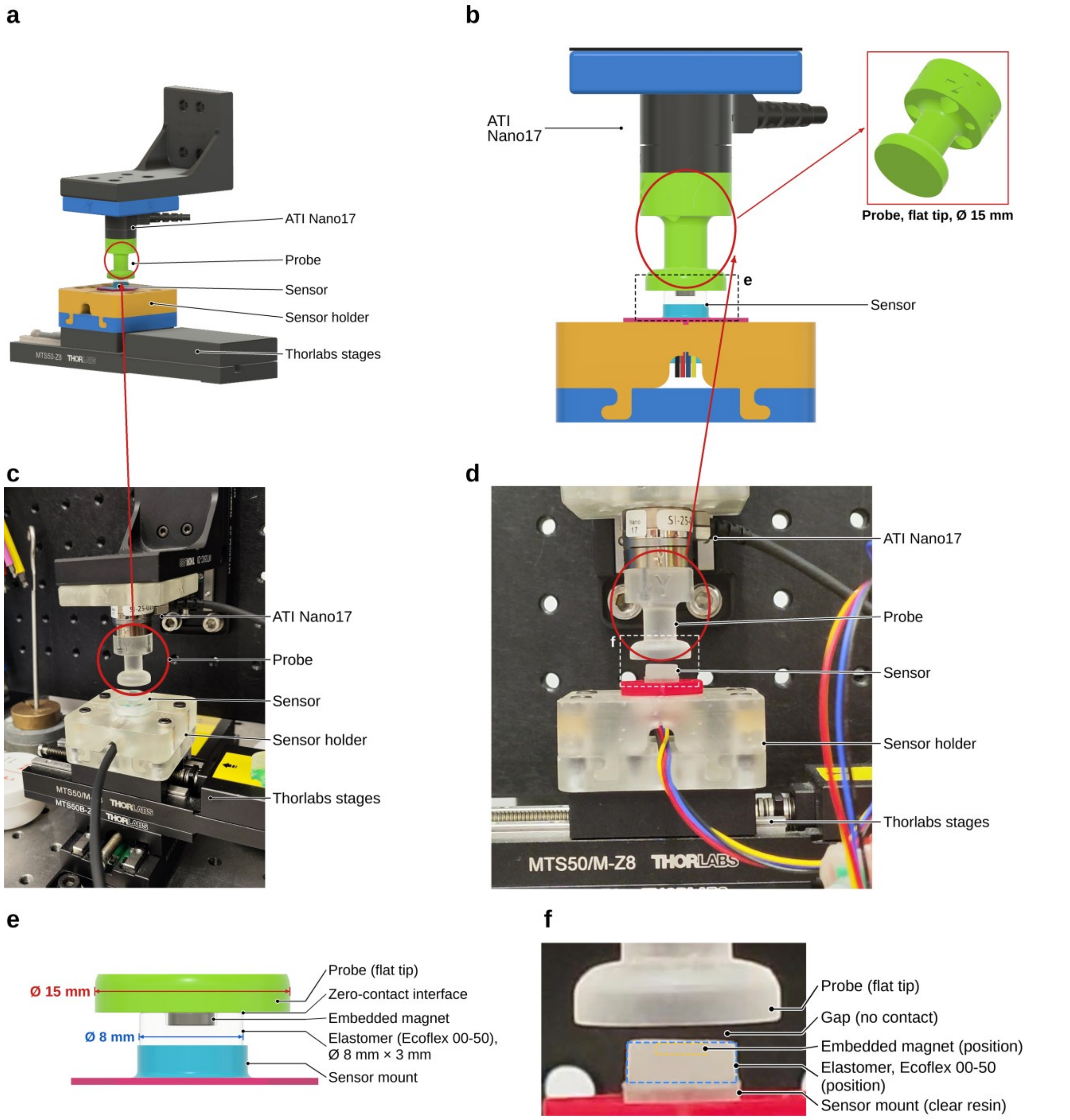


**Figure S8.** Calibration rig. (a) CAD model: ATI Nano17 transducer with probe (circled) above the sensor holder on motorized stages. (b) Section through probe and sensor; inset, 15-mm flat probe tip. (c) Rig photograph (earlier holder version). (d) Probe above a sensor during calibration (Video S1). (e, f) Probe–elastomer interface at zero contact in CAD and photograph; dashed outlines mark the embedded elastomer and magnet.

#### S6.4 Calibration Processing and Force Equations

Calibration data were processed by Sensor Tracking Studio in three stages, each of which saves its results to a file. In stage 1, every sample is labeled with its displacement step (the *z*-stage encoder position snapped to a 0.1-mm grid; transient segments shorter than 25 samples are merged with the adjacent step), cycle (1–5) and phase (loading or unloading). In stage 2, the minimum, maximum, median and mean of the reference force, indentation and raw $B_z$ are computed for every step. In stage 3, polynomials are fitted to the step medians by unweighted least-squares regression (scikit-learn), separately for each cycle and for all cycles combined, for the loading and unloading branches, and for force and indentation as targets, at both second and third order:

$$F_N = c_3 B_z^3 + c_2 B_z^2 + c_1 B_z + c_0 \quad \text{(with } c_3 = 0 \text{ for the second-order fit).}$$

No temporal filter was applied to the calibration data. The force readings in this study used, for each sensor, the cubic equation fitted to the loading branch of cycle 3 (19 step medians per curve; Figure S9, Table S4). The cubic was chosen because the second-order fit systematically over-predicted low forces (Figure S10): below 40 gf of reference force, the median bias across the twelve sensors was +5.0 gf for the second-order and +0.3 gf for the cubic fit, and the median root-mean-square error (RMSE) over the whole calibrated range was 7.1 gf and 1.9 gf, respectively. With the second-order fit, this bias gave false readings below 1 gf at sites without contact. Loading–unloading hysteresis (Figure S11), quantified from the force difference at matched displacement steps relative to the full-scale range, averaged 7.2–9.4% of full scale (maximum 11.1–15.5%) and was larger than the fit error; for this reason a single (loading) branch was applied consistently to all sensors and runs. Cycle-to-cycle repeatability was quantified as the standard deviation of the step-median force across the five loading cycles at matched displacement.

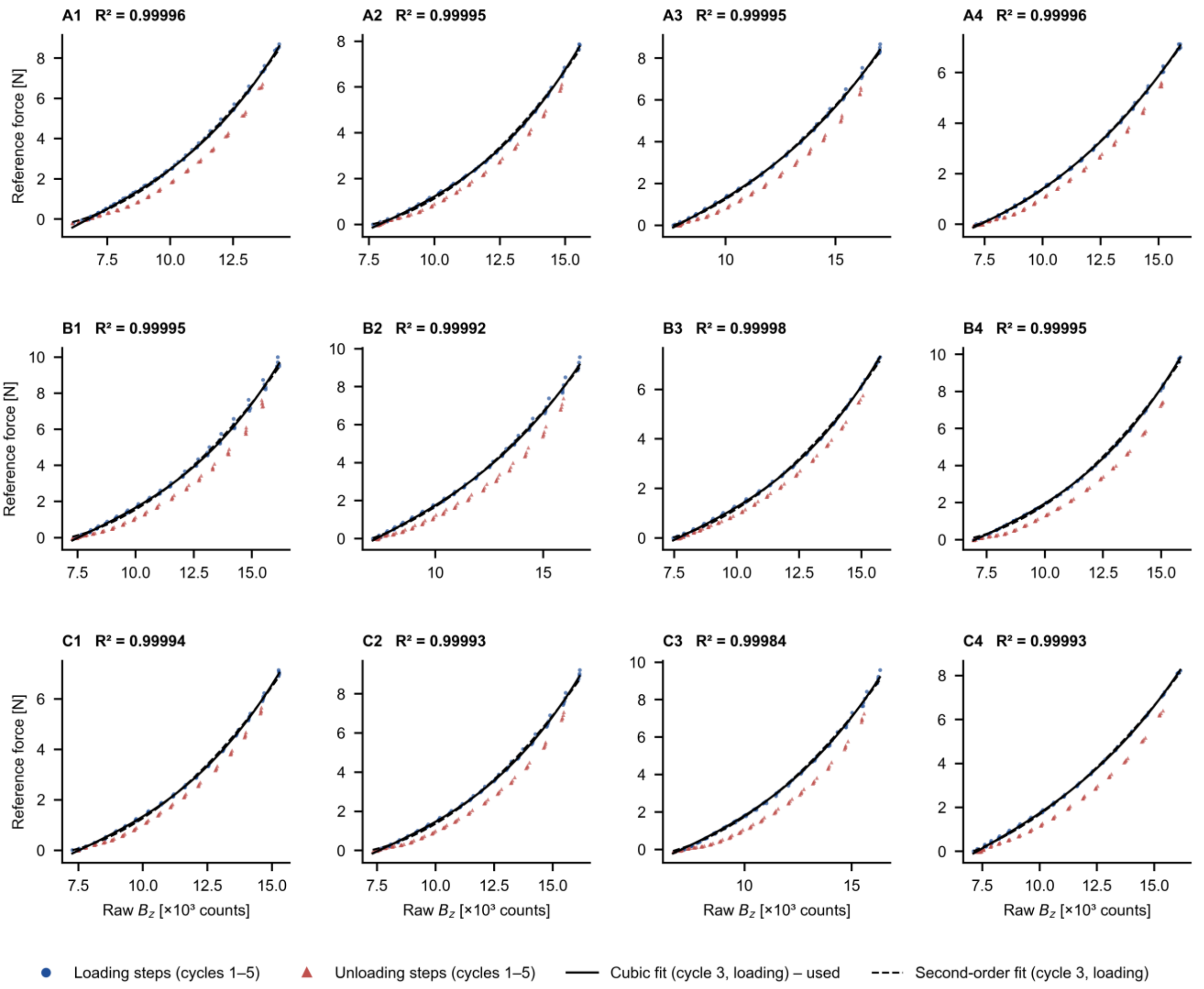


**Figure S9.** Calibration data of the twelve sensors: step-median reference force against raw $B_z$ (loading, circles; unloading, triangles), with cubic (solid) and second-order (dashed) fits to cycle-3 loading; $R^2$, cubic fit.

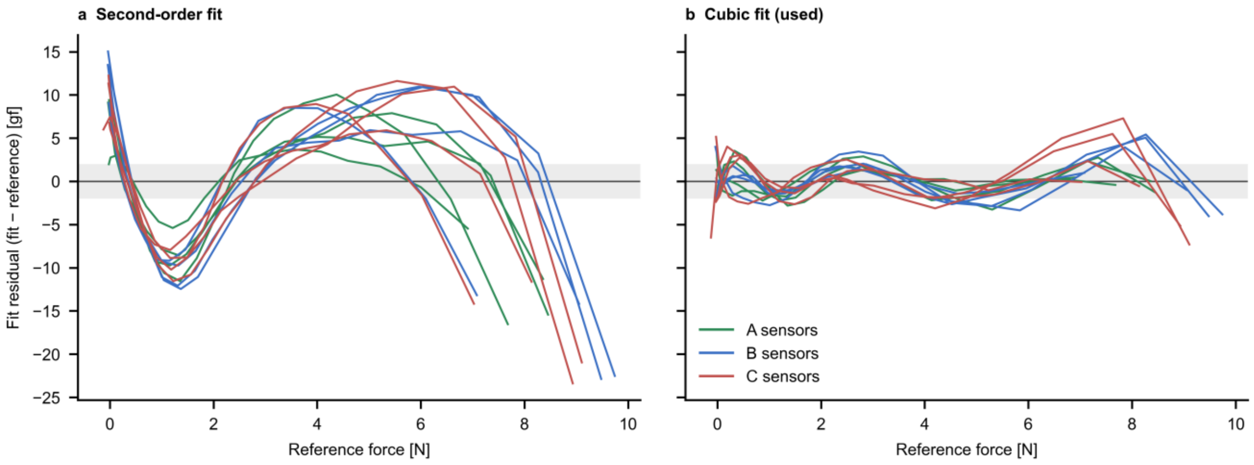


**Figure S10.** Residuals of (a) second-order and (b) cubic fits; gray band, ±2 gf.

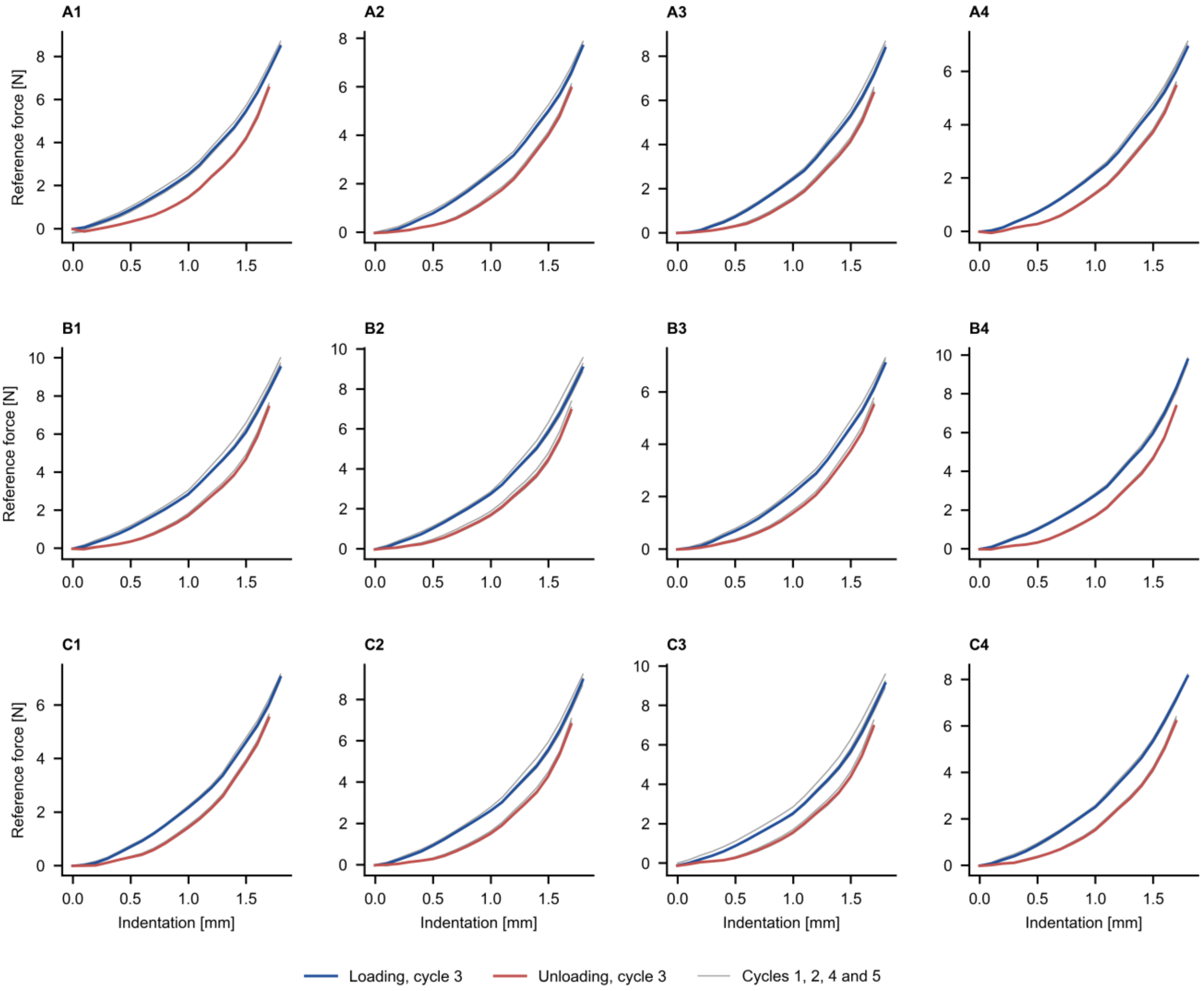


**Figure S11.** Loading–unloading hysteresis of all twelve sensors. Step-median reference force against indentation for loading (blue) and unloading (red); cycle 3 is drawn bold and cycles 1, 2, 4 and 5 thin, showing the cycle-to-cycle repeatability.

**Table S4.** Cubic calibration equations (loading branch, cycle 3) and calibration performance (set S1_MatC). $F_N$ in N and $B_z$ in raw counts; hysteresis, mean (maximum) % of full scale; repeatability, mean (maximum) SD of step-median force over five cycles.

| Sensor | $c_3$ | $c_2$ | $c_1$ | $c_0$ | Range [N] | $R^2$ | RMSE [N] | Hysteresis [% FS] | Repeatability [N] |
|---|---|---|---|---|---|---|---|---|---|
| **A1** | $7.277\times10^{-12}$ | $-1.409\times10^{-7}$ | $1.570\times10^{-3}$ | -6.416 | 0–8.70 | 0.99996 | 0.015 | 9.4 (15.0) | 0.106 (0.147) |
| **A2** | $7.517\times10^{-12}$ | $-1.730\times10^{-7}$ | $1.861\times10^{-3}$ | -7.617 | 0–7.87 | 0.99995 | 0.015 | 8.7 (13.2) | 0.082 (0.161) |
| **A3** | $3.105\times10^{-12}$ | $-6.333\times10^{-8}$ | $9.822\times10^{-4}$ | -5.297 | 0–8.67 | 0.99995 | 0.018 | 8.0 (13.8) | 0.076 (0.191) |
| **A4** | $1.974\times10^{-12}$ | $-1.544\times10^{-8}$ | $3.418\times10^{-4}$ | -2.459 | 0–7.13 | 0.99996 | 0.013 | 8.0 (12.7) | 0.052 (0.119) |
| **B1** | $6.779\times10^{-12}$ | $-1.563\times10^{-7}$ | $1.840\times10^{-3}$ | -7.896 | 0–10.00 | 0.99995 | 0.020 | 9.3 (15.1) | 0.116 (0.234) |
| **B2** | $3.605\times10^{-12}$ | $-7.270\times10^{-8}$ | $1.083\times10^{-3}$ | -5.432 | 0–9.56 | 0.99992 | 0.024 | 9.2 (15.5) | 0.121 (0.329) |
| **B3** | $5.879\times10^{-12}$ | $-1.319\times10^{-7}$ | $1.478\times10^{-3}$ | -6.233 | 0–7.30 | 0.99998 | 0.009 | 7.5 (12.5) | 0.074 (0.154) |
| **B4** | $6.030\times10^{-12}$ | $-1.187\times10^{-7}$ | $1.350\times10^{-3}$ | -5.716 | 0–9.85 | 0.99995 | 0.020 | 8.5 (13.4) | 0.063 (0.136) |
| **C1** | $6.212\times10^{-12}$ | $-1.349\times10^{-7}$ | $1.464\times10^{-3}$ | -6.035 | 0–7.15 | 0.99994 | 0.016 | 7.2 (11.1) | 0.051 (0.122) |
| **C2** | $6.283\times10^{-12}$ | $-1.416\times10^{-7}$ | $1.631\times10^{-3}$ | -6.972 | 0–9.22 | 0.99993 | 0.023 | 9.1 (14.2) | 0.110 (0.244) |
| **C3** | $3.689\times10^{-12}$ | $-6.312\times10^{-8}$ | $8.776\times10^{-4}$ | -4.365 | 0–9.58 | 0.99984 | 0.034 | 8.3 (13.7) | 0.178 (0.325) |
| **C4** | $3.696\times10^{-12}$ | $-7.549\times10^{-8}$ | $1.107\times10^{-3}$ | -5.495 | 0–8.23 | 0.99993 | 0.020 | 9.2 (15.1) | 0.052 (0.083) |

#### S6.5 Data Acquisition and Processing

During the force experiments, the twelve-sensor stream was transmitted by the Teensy 4.1 over a 115 200-baud USB serial link. Sensor Tracking Studio polled the serial buffer every 10 ms (nominal 100-Hz software polling) and read up to 50 available lines per poll; the rate at which the firmware sent data, and therefore the actual sampling rate of each sensor, did not depend on this polling interval. No digital low-pass, moving-average or other temporal filter was applied. Raw $B_z$ values were converted sample by sample using the sensor-specific cubic equation. Before each run, Phantom 3 was prepared at the 77-day (704-mL) baseline and, with the helmet not yet fitted and all sensors unloaded, the operator activated TARE ALL; the force predicted from the current raw value was stored as a sensor-specific offset and subtracted from all subsequent readings (a negative predicted value was stored as zero). Negative readings were set to zero. No automatic baseline tracking, temperature compensation or drift correction was implemented.

Force was calculated in newtons and converted to gram-force for display and reporting using 1 N = 101.97 gf (1 gf = 9.80665 mN). Each session log contains one row per socket per frame, with a computer-clock timestamp (millisecond precision, assigned on serial arrival), the sensor identity, the socket, the operator-entered volume label, raw $B_z$, force in newtons and gram-force and the equivalent indentation depth. The twelve channels followed the serial scan order but did not share a hardware-generated timestamp; malformed lines were discarded and packet loss was not separately counted.

The final 704-mL segment, recorded after volume release, was evaluated in the same way as the other volume steps (Section 5.6, main text) and is reported as "704 mL (after release)".

#### S7. Complete Force Results (n = 7)

Force experiments comprised seven fittings of the instrumented helmet on Phantom 3, labeled Fittings 1–7 in chronological order; Fitting 3 is the representative run shown in Figure 5a (main text). The continuous traces of all seven fittings, including the 212-day (900-mL) extension and volume release, are shown in Figure S12 and in Video S3 for Fitting 6. Table S5 lists the per-sensor forces at every volume step, Table S6 gives the detailed values at 175 days (860 mL), and Table S7 gives the regional load shares.

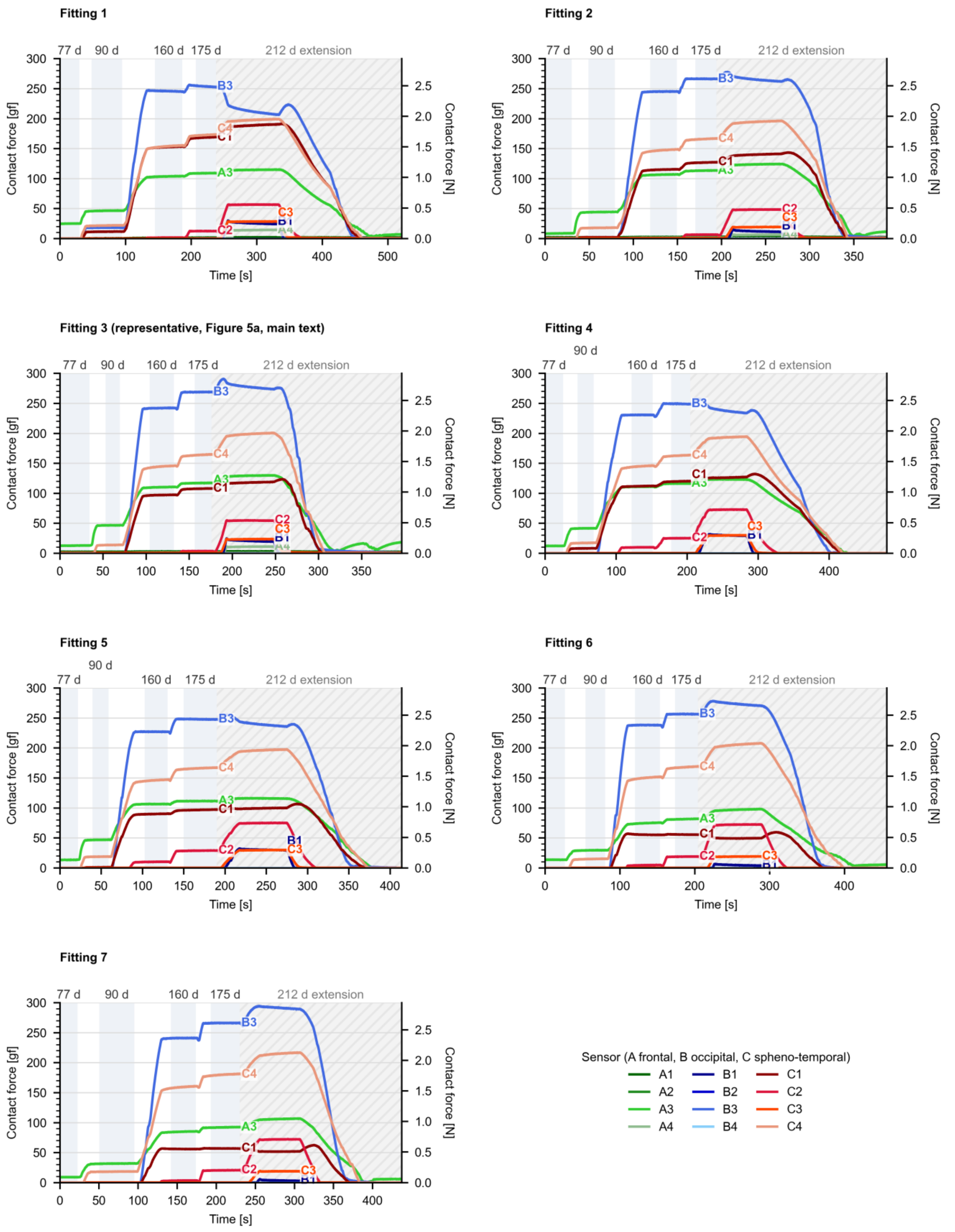


**Figure S12.** Continuous force traces of all seven fittings, plotted as in Figure 5a (main text); hatched region, 212-day (900-mL) extension and volume release.

**Table S5.** Per-sensor contact force [gf] at each volume step, mean ± standard deviation across seven fittings (n = 7). “< 0.5” indicates no measurable force (all fittings below 0.5 gf). Columns shaded in amber belong to the main study interval.

| Sensor | 77 d (704 mL) | 90 d (732 mL) | 160 d (843 mL) | 175 d (860 mL) | 212 d (900 mL) | After release (704 mL) |
|---|---|---|---|---|---|---|
| **A1** | 0.9 ± 1.3 | 0.9 ± 1.3 | 1.2 ± 1.6 | 1.3 ± 1.6 | 1.4 ± 1.8 | 1.2 ± 1.5 |
| **A2** | < 0.5 | < 0.5 | < 0.5 | < 0.5 | < 0.5 | < 0.5 |
| **A3** | 13.3 ± 5.4 | 40.8 ± 7.3 | 99.5 ± 14.1 | 105.7 ± 13.6 | 115.6 ± 11.1 | 5.8 ± 4.8 |
| **A4** | < 0.5 | < 0.5 | 0.3 ± 0.5 | 0.4 ± 0.5 | 4.4 ± 6.0 | < 0.5 |
| **B1** | < 0.5 | < 0.5 | < 0.5 | < 0.5 | 17.7 ± 11.4 | < 0.5 |
| **B2** | < 0.5 | < 0.5 | < 0.5 | < 0.5 | < 0.5 | < 0.5 |
| **B3** | < 0.5 | 2.7 ± 6.7 | 238.5 ± 7.2 | 258.3 ± 8.7 | 255.8 ± 27.8 | 0.5 ± 0.8 |
| **B4** | < 0.5 | < 0.5 | < 0.5 | < 0.5 | < 0.5 | < 0.5 |
| **C1** | < 0.5 | 3.1 ± 4.6 | 96.7 ± 34.4 | 104.5 ± 39.9 | 110.5 ± 49.4 | < 0.5 |
| **C2** | < 0.5 | < 0.5 | 4.1 ± 4.2 | 16.3 ± 9.4 | 64.3 ± 10.9 | < 0.5 |
| **C3** | < 0.5 | < 0.5 | < 0.5 | < 0.5 | 23.9 ± 5.2 | 0.2 ± 0.3 |
| **C4** | < 0.5 | 17.3 ± 2.6 | 149.4 ± 5.4 | 168.5 ± 5.9 | 200.4 ± 7.7 | < 0.5 |

**Table S6.** Contact force at 175 days (860 mL) for the sensors that carried measurable load, across seven fittings (n = 7). Nominal pressure as defined in Section 5.6 (main text); CV, coefficient of variation.

| Sensor | Region | Mean ± SD [gf] | Mean ± SD [N] | Range [gf] | CV [%] | Nominal pressure [kPa] |
|---|---|---|---|---|---|---|
| **A3** | Frontal | 105.7 ± 13.6 | 1.04 ± 0.13 | 81.3–117.0 | 13 | 20.6 |
| **B3** | Occipital | 258.3 ± 8.7 | 2.53 ± 0.09 | 247.8–268.8 | 3 | 50.4 |
| **C1** | Spheno-temporal (left) | 104.5 ± 39.9 | 1.02 ± 0.39 | 55.5–168.5 | 38 | 20.4 |
| **C2** | Spheno-temporal (left) | 16.3 ± 9.4 | 0.16 ± 0.09 | 3.3–28.6 | 58 | 3.2 |
| **C4** | Spheno-temporal (right) | 168.5 ± 5.9 | 1.65 ± 0.06 | 162.8–180.0 | 4 | 32.8 |

**Table S7.** Regional load shares and summed sensor load at each volume step (n = 7). Values are the mean (minimum–maximum) across fittings; shares are defined in Section 5.6 (main text). Shares are not given after release because the summed load was close to zero.

| Volume step | Frontal (A1–A4) [%] | Occipital (B1–B4) [%] | Spheno-temporal (C1–C4) [%] | Summed load [gf] |
|---|---|---|---|---|
| **77 d (704 mL)** | 99 (97–100) | 0 (0–0) | 1 (0–3) | 14 (9–25) |
| **90 d (732 mL)** | 65 (48–78) | 3 (0–18) | 32 (22–38) | 65 (44–98) |
| **160 d (843 mL)** | 17 (14–19) | 41 (37–46) | 42 (40–47) | 590 (523–659) |
| **175 d (860 mL)** | 16 (14–18) | 40 (35–44) | 44 (41–49) | 655 (580–718) |
| **212 d (900 mL)** | 15 (13–17) | 35 (28–39) | 50 (47–56) | 794 (720–839) |
| **After release (704 mL)** | – | – | – | 8 (0–18) |

**Extension to 212 days (900 mL).** Beyond the Helmet 1 interval, two further sites came into contact: the occipital site B1 (17.7 ± 11.4 gf) and the right spheno-temporal site C3 (23.9 ± 5.2 gf). Force increased further at C4 (200.4 ± 7.7 gf) and C2 (64.3 ± 10.9 gf), whereas B3 did not increase further on average (255.8 ± 27.8 gf) and became more variable between fittings (211–291 gf); in Fitting 1, B3 force decreased during the extension while C1 and C4 increased (Figure S12). The spheno-temporal share rose to about half of the summed sensor load (Table S7). Sites A2, B2 and B4 recorded no measurable force at any volume.

**Volume release.** On return to 704 mL, forces at all sensing sites fell back to the baseline range: A3 recorded 5.8 ± 4.8 gf (loading baseline 13.3 ± 5.4 gf) and all other sites less than 1.5 gf on average, showing that the zeroed sensors returned to their starting values after loading.

#### S8. Supplementary Imaging Material

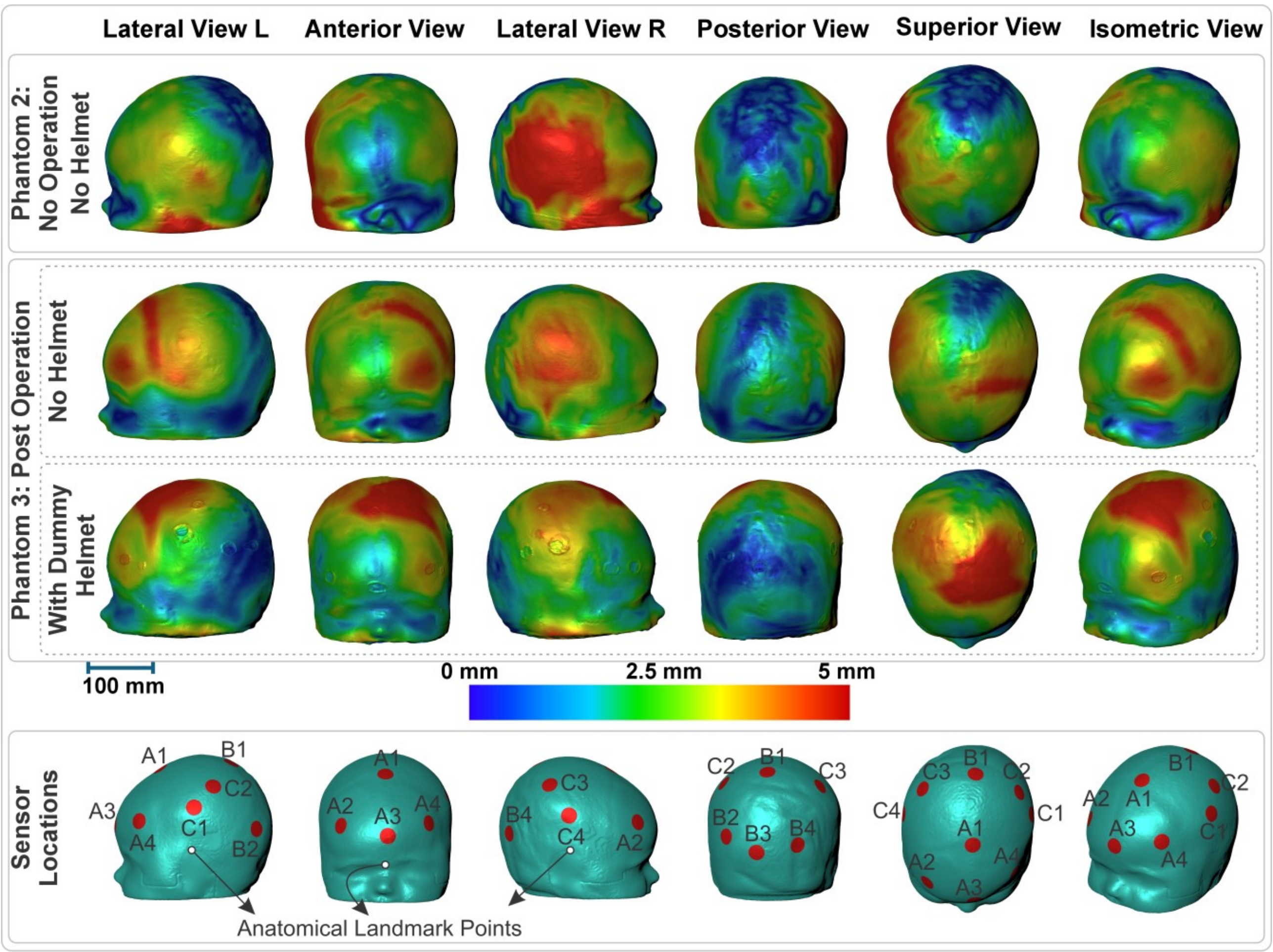


**Figure S13.** Surface displacement from 77 to 212 days (704 to 900 mL) for the three configurations of Figure 6 (main text), in six views; scale bar, color scale and sensor markers as in Figure 6.

**Sensor projection.** The approximate sensor positions shown in Figure 6 (main text) and Figure S13 were projected onto the CT-derived skin surface from the helmet CAD and by visual correspondence with the sensor openings of the dummy helmet.

**Section views.** The section overlays of Figure 7 (main text) were made by exporting the CT-derived helmet, 77-day (704-mL) and 175-day (860-mL) skin surfaces from Avizo 2022.1 (Thermo Fisher Scientific, USA), aligning them in a common coordinate system and sectioning them in Fusion 360 (Autodesk, USA) along the P1 sagittal, P2 and P3 coronal and P4 horizontal planes.

**Longitudinal validation.** Phantom 1, manufactured at 572 mL from the patient’s 56-day CT (Table S3), was expanded *in vitro* to the 77-day volume (704 mL), and its CT-derived external skin surface was compared with the patient’s 76-day pre-operative surface scan (Figure S14). On the common color scale, most of the cranial vault fell within 0–2.5 mm, with only limited regions approaching 5 mm. The prominent difference along the lower edge coincided with missing coverage in the clinical surface scan and was treated as an acquisition gap rather than a true disagreement between the surfaces. Because Phantom 1 was built from the patient’s CT rather than from the companion finite-element model, this comparison tests the combined fabrication-and-expansion process, but it does not separate manufacturing from expansion accuracy.

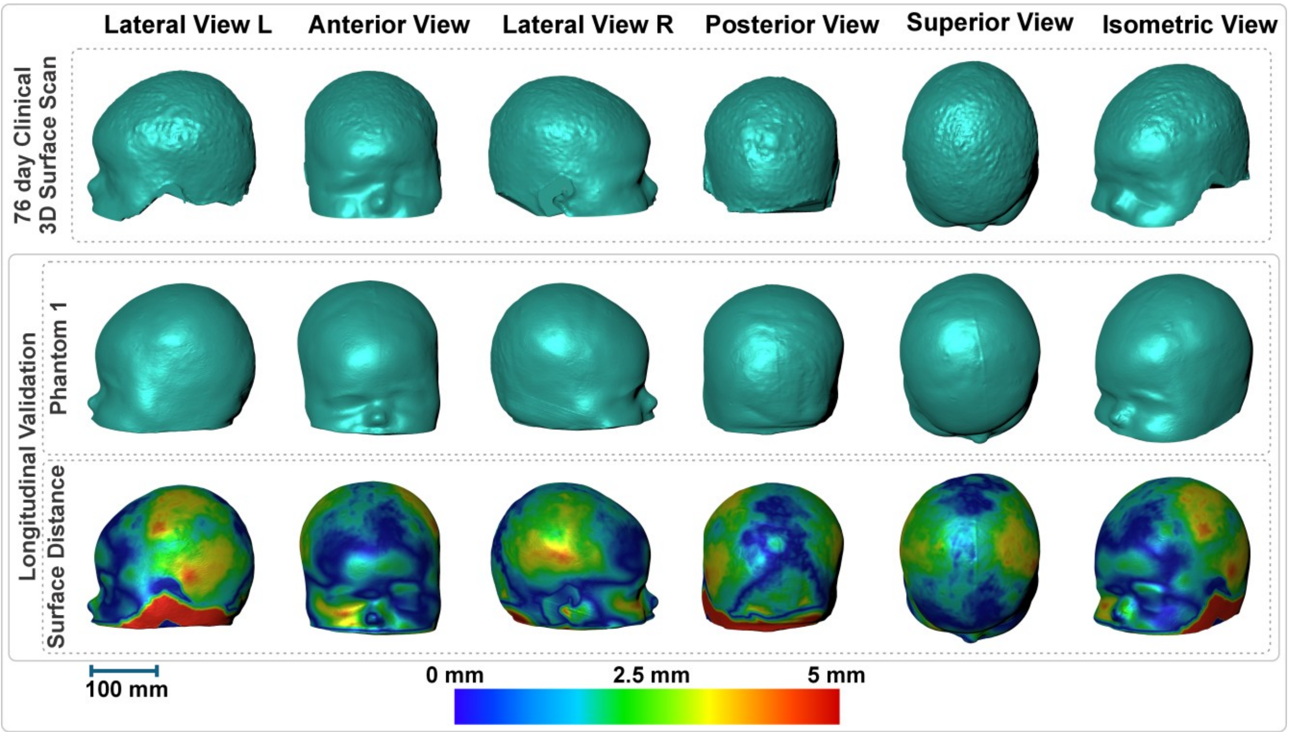


**Figure S14.** Longitudinal validation of Phantom 1. Top: the patient's 76-day pre-operative surface scan. Middle: Phantom 1 after *in vitro* expansion from 56 to 77 days (572 to 704 mL). Bottom: surface-distance map between the two surfaces; scale bar and 0–5-mm color scale as in Figure 4 (main text).

### S9. Supplementary Videos

| Video | Content |
|---|---|
| **Video S1** | Sensor calibration: one loading and unloading cycle of a sample sensor of the same design on the calibration rig, with automatic contact detection, 0.1-mm steps to 1.8 mm and 10-s holds (9× speed). |
| **Video S2** | Preparation before Fitting 6: serial connection, selection of the cubic equation, calibration cycle and sensor set, tare, finger check of each sensor, helmet fitting and mounting in the test rig (real time; helmet fitting and final checks at 8× speed). |
| **Video S3** | Force run of Fitting 6: expansion from 77 to 212 days (704 to 900 mL) and release to 704 mL, with the live Sensor Tracking Studio dashboard and three camera views (16× speed). |

### S10. Code and Data Availability

**Helmet contact-force analysis** [10]: force logs of the seven fittings (Fittings 1–7) recorded on 24 September 2026; the calibration step medians of the twelve sensors (set S1_MatC); and the scripts that generate Figure 5 (main text), Figures S9–S12 and Table S4. Code under the PolyForm Noncommercial License 1.0.0; data and images under CC BY-NC 4.0.

**Sensor Tracking Studio** [9]: source code of the application described in Section S6.2, including the calibration pipeline, the equation set used in this study and the Teensy 4.1 firmware that streams the twelve sensors. Code under GPL-3.0; example datasets and reports under CC BY-NC 4.0.

**Tactile-sensor calibration rig** [11]: control and acquisition software of the rig described in Section S6.3, including its firmware. Version 1.0 (February 2026) recorded the calibrations used in this study; later versions keep the same stream-file format. Code under GPL-3.0.

The raw calibration recordings (about 2.2 GB) are available from the corresponding author upon reasonable request.
